# Thalamic NRXN1-Mediated Input to Human Cortical Progenitors Drives Upper Layer Neurogenesis

**DOI:** 10.1101/2025.04.25.650717

**Authors:** Claudia V. Nguyen, Antoni Martija, Patricia R. Nano, Jose A. Soto, Daniel C. Jaklic, Jessenya Mil, Rista White, Jacqueline M. Martin, Dakshesh Rana, Daniel H. Geschwind, Aparna Bhaduri

**Affiliations:** Department of Biological Chemistry, David Geffen School of Medicine, University of California, Los Angeles, CA, USA; Department of Human Genetics, David Geffen School of Medicine, University of California, Los Angeles, CA, USA; Department of Psychiatry and Biobehavioral Sciences, David Geffen School of Medicine, University of California, Los Angeles, CA, USA; Department of Neurology, David Geffen School of Medicine, University of California, Los Angeles, CA, USA

## Abstract

According to the protocortex hypothesis, extrinsic thalamic signaling is necessary for refining cortical areas and cell types, but the mechanism by which these inputs shape the development and expansion of the human cortex remains largely unexplored. We fuse cortical and thalamic organoids to study this process. Using single-nuclei RNA-sequencing and cellular imaging, we discover that thalamic signals during a critical period promote human cortical upper-layer neurogenesis. In assembloid models and human primary cortex, we find NRXN1 mediates thalamic axon contact with primate-enriched outer radial glia, driving developmental gene expression changes. Genetic perturbation of NRXN1 in thalamic neurons reduces these contacts and attenuates cortical upper-layer neurogenesis. These findings in human developmental models suggest a novel role for thalamic regulation of primate outer radial glia cell fate.

## Introduction

The unique cognitive features of the human cortex can be largely attributed to the evolutionary expansion of the neocortex compared to other mammals; this expansion emerges during prenatal development. During development, signals both intrinsic to the cortex and extrinsically, from neighboring brain areas, influence the trajectory of cortical cell types. The primary source of developmental extrinsic cues to the cortex is the thalamus, which serves as a relay center for sensory information. During development, the thalamus engages in coordinated activity with the cortex, including via the extension of reciprocal axonal afferents to and from the cortex (*1–6*). Previous work in rodent models has demonstrated that thalamocortical afferents (TCAs) that project from the thalamus to the cortex play an integral role in cortical layer and areal fate refinement, as well as in neuronal migration and maturation (*7–10*). Longstanding questions in the study of cortical arealization have explored the relative merits of a protomap model whereby cortical radial glia are predetermined to a specific fate (*11*), versus a protocortex where the cortex develops agnostic to areal identity until extrinsic cues deliver key patterning input (*12*). While the consensus in the field has settled on a mix of these models (*13*), similar reconciliation of how thalamic signals regulate cues during cortical neurogenesis are more challenging as coordinated development and reciprocal interaction between the cortex and thalamus are difficult to disentangle *in vivo* (*6*).

There are several unique properties of human cortical development that differentiate it from traditional rodent model organisms. Human brain development, and that of other gyrencephalic mammals, differs especially in its expanded pool of complex progenitor subtypes called radial glia, which are responsible for deciphering these intrinsic and extrinsic cues that ultimately manifest in mature cell types and contribute to cortical expansion (*14–16*). In comparison to rodents, primates have an expanded population of progenitors, called outer radial glia (oRG), that comprise the majority of cells present in the outer subventricular zone (OSVZ) of the developing cortex (*14, 17*). These and other radial glia subtypes identified in the developing human brain give rise to an expanded layer 2/3 in humans and nonhuman primates compared to rodents (*18*). In concert with these unique features of cortical development, TCAs also have both divergent and conserved characteristics between species. Notably, TCA innervation occurs much earlier in primate development than in rodents. Thalamic afferents extend toward the mouse and rat neocortex at embryonic day E12/13 and enter via the intermediate zone by E15/16 (*1–3, 19*). In primates, neocortical innervation by TCAs occurs immediately after neuroepithelial stages (*20–22*). Interestingly, a well-conserved feature of rodent and nonhuman primate development is that TCAs exhibit transient synapses with the cortical subplate (*22–29*). Taken together, the earlier entry of TCAs and their known transient roles in the subplate prompted us to question whether TCAs may engage in other transient signaling roles, including with primate-enriched oRG progenitors.

Across mammalian systems, questions remain about the specific role that TCAs play in orchestrating cortical cell fate decisions, especially regarding the mechanism by which they influence cortical differentiation and arealization. An even more limited understanding exists about how or why TCAs communicate with cortical radial glia during human prenatal development, though at postnatal periods aberrant TCA connectivity has been associated with neuropsychiatric and neurodevelopmental disorders (*30–33*). Thus, we sought to explore what role TCAs play in driving human cortical cell fate specification and the ways in which these axons communicate with cortical progenitors.

We chose to leverage a previously described *in vitro* stem cell-derived model of the human cortex and thalamus, called a thalamocortical (TC) assembloid (*34*), to determine the unique contributions of thalamic input to cortical cell fate specification. Landmark studies in the field of neurodevelopment have previously established protocols generating organoids resembling human brain regions, such as the cortex, that have enabled us to model key features of human brain development (*35, 36*). The further innovation of the assembloid, a culture which fuses two or more organoid types (*37*), has expanded our ability to model connectivity between brain regions during development. Park and colleagues have generated TC assembloids (*34*), adaptations of which have since been used to model aspects of neurodevelopmental disorders characterized by thalamocortical dysregulation (*38–40*). These assembloid systems uniquely afford the ability to not only study human development and primate-enriched cell types such as oRGs, but also provide an opportunity to systematically manipulate the addition or removal of thalamic input to the developing cortex at timepoints of interest. This experimental accessibility allows for the interrogation of how thalamic inputs regulate human cortical progenitors.

Our study identifies neurexin 1 (NRXN1) as a facilitator of TCA to oRG communication. Neurexins are mainly characterized for their role as membrane cell-adhesion molecules that bind to neuroligins (NLGNs). In the brain, neurexins are primarily known for their roles in synaptic development and stabilization (*41–48*), though in other tissues and contexts they have been shown to be involved in cellular adhesion and filopodial extension and branching (*49–54*). Here, we provide evidence for a non-canonical role of neurexins, whereby thalamic NRXN1 mediates a physical interaction with cortical progenitors. We demonstrate that this interaction is necessary for the expansion of cortical upper layer neurogenesis, thus extending our ability to understand the thalamus’s role in promoting areal fate to additional roles in regulating human progenitor cell fate choices.

## Results

### Thalamic input increases the proportion of cortical excitatory neurons

There remain many open questions regarding the role of thalamic afferents that innervate the cortex during human prenatal development. To address this gap, we leveraged the assembloid system to investigate foundational aspects of how TCAs impact cortical fate specification and maturation. We implemented previously described protocols for generating cortical organoids (*55–57*), and adapted a previously published thalamic organoid protocol (*34, 58*). We further optimized the thalamocortical fusion protocol (*34*), fusing them at week 3 in a media containing half thalamic and half cortical media (Fig S1A). These assembloids better retain their individual regional identities over the course of differentiation as assessed using thalamic and cortical markers (Fig S1B, Fig S2). Corticocortical assembloids grown in the same media mix as our thalamocortical (TC) assembloids were used as controls for phenotypes arising from media signals or the fusion process itself (Fig 1A, Fig S1C).

**Figure 1.**
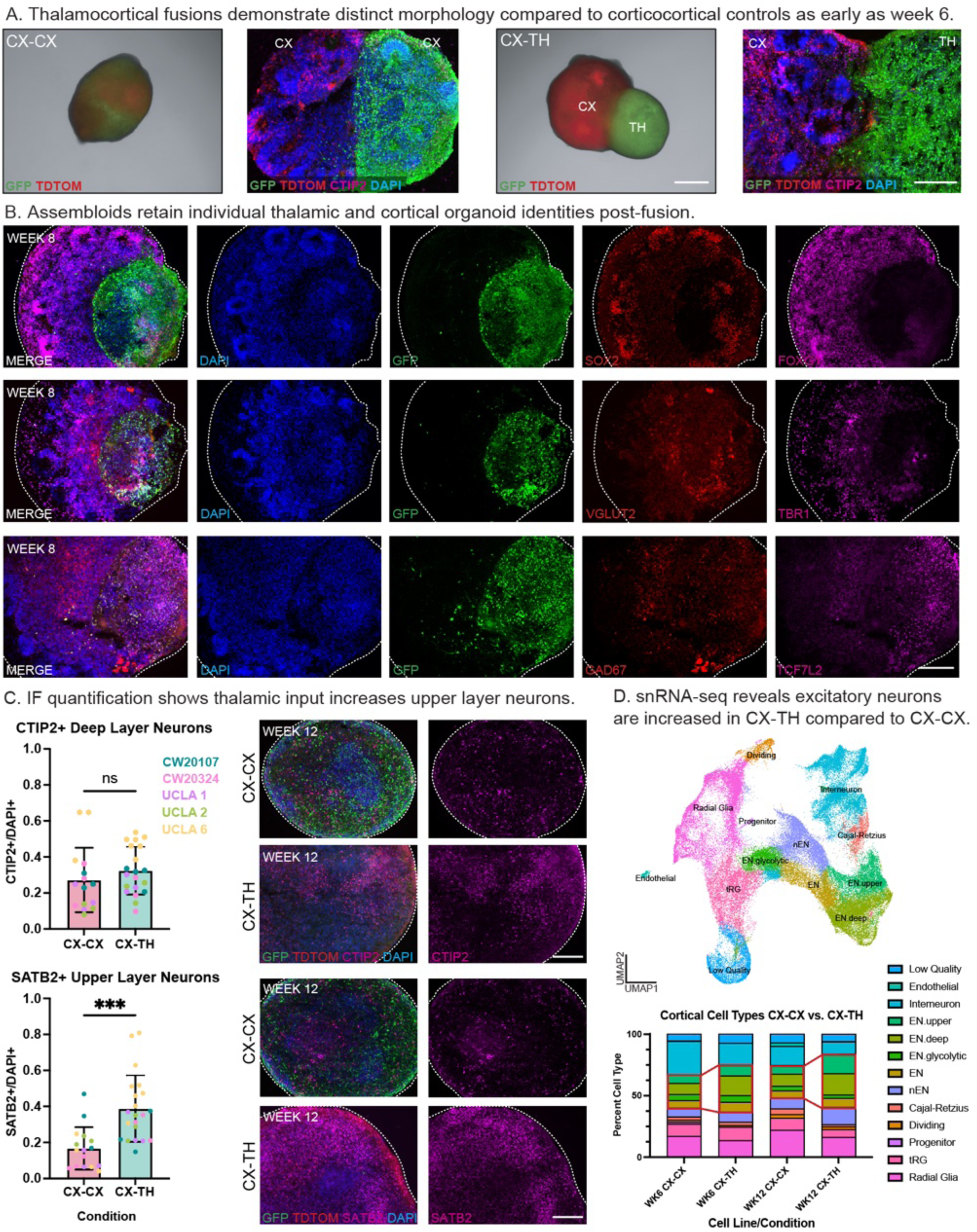
Comparisons of thalamocortical (CX-TH) assembloids to corticocortical (CX-CX) controls reveals fusion with thalamus increases proportion of cortical excitatory neurons. A) Thalamocortical (TC) assembloids were used to study how the thalamus influences cortical cell fate. Assembloids were generated at week 3, and pictured here live at week 6 (1st & 3rd image) and were also immunostained for further analysis. Corticocortical assembloids were used as a control (CX-CX, left) and demonstrate a difference in morphology compared to thalamocortical (CX-TH, right) counterparts. Prior to aggregation, stem cells were lentivirally labeled with LV-EF1a-tdTomato (cortical side) or LV-EF1a-GFP (thalamic side) to trace reciprocal afferents by counterstaining for these markers in immunofluorescent staining (2nd & 4th image). Staining for CTIP2 (magenta) was further used to confirm the identity of the cortical half of the assembloid. Scale bar = 500 μm. B) Canonical markers for progenitors as well as mature cortical and thalamic markers were used to confirm distinct identities were preserved on either side of the TC assembloids. GFP was used to identify the thalamic cells in the assembloid. Areas of the assembloid positive for GFP were positive for thalamic-enriched markers VGLUT2 (glutamatergic), GAD67 (GABAergic), and TCF7L2 (pan-thalamic), as expected. Conversely, GFP-cortical cells were enriched for cortex-specific markers such as FOXG1 (CX progenitors) and TBR1 (cortical layer 6). Both thalamus and cortex contained cells positive for radial glia marker SOX2, as predicted. Representative images are from week 8 assembloids generated from UCLA 2. Analysis was performed in 5 different stem cell lines and for each line, 3 - 4 technical replicates were examined. Scale bar = 250 μm. C) Proportions of CTIP2+ (top) and SATB2+ (bottom) cells across 5 lines were quantified and shown in histogram as a ratio over total DAPI+ cells to determine the proportion of deep and upper layer neurons, respectively. Comparisons of these ratios demonstrate a significant increase in the proportion of upper layer neurons (SATB2+, p = 0.0002, t-test) in CX-TH assembloids compared to CX-CX controls. No significant change in the proportion of deep layer neurons (CTIP2+) was observed. Technical replicates of 3 - 9 organoids from each of 5 lines were quantified to achieve per condition averages (left). Representative images (right) are from week 12 assembloids generated from UCLA 6. Scale bar = 250 μm. D) Quantification of relative cell types from single-nuclei RNA-sequencing data shows an increase in excitatory neuron specification in CX-TH compared to CX-CX controls across 5 cell lines. UMAP (top) depicts annotated clusters from analysis of week 6 and 12 CX-TH and CX-CX single-nuclei data. Bar graph (bottom) depicts cell type percentages from each of these time points and conditions, as extracted from the UMAP. Red boxes highlight excitatory neuron groups to demonstrate increases in these populations with the addition of thalamus to assembloids.

Cell types in TC assembloids (CX-TH) versus corticocortical control assembloids (CX-CX) were initially explored via immunostaining and quantification. We stained for CTIP2 (gene *BCL11B*) and SATB2, markers of deep layer and upper layer cortical neurons, respectively. We quantified these markers in the cortical regions of week 12 assembloids, a time point that corresponds with the end of neurogenesis. We identified a 2.3-fold increase in SATB2 in CX-TH compared to CX-CX across 5 stem cell lines, spanning male, female, human embryonic stem cell (hESC) and induced pluripotent stem cell (hiPSC) backgrounds (Fig 1C, Table S1). However, no significant difference was observed in CTIP2 between conditions (Fig 1C). Increases in upper layer neuronal specification corroborate the role of TCAs in promoting superficial layer expansion previously shown in rodent models (*10, 59*). These studies, however, showed a reciprocal decrease in deep layer fate that was not observed in our study.

Additionally, single-nuclei RNA-sequencing (snRNA-seq) was performed in week 6 and 12 assembloid samples across the same 5 cell lines (Fig S3A - B). Cell type annotation and quantification corroborated increases in excitatory neuron specification on the cortical side of the CX-TH condition as compared to CX-CX (Fig 1D). The snRNA-seq data demonstrates increases in excitatory cell populations in the CX-TH assembloids at weeks 6 and 12, suggesting that transcriptional differences may precede protein-level phenotypes. These same phenotypes were observed across individual cell lines as well (Fig S3C).

### Thalamic input in early neurogenesis is necessary and sufficient to expand cortical upper layer specification

To explore when the protein-level phenotypes emerge, we quantified cell types via immunofluorescence in cortical regions across assembloid conditions. Deep layer populations (CTIP2+) remained consistent between groups across timepoints, while the significant difference in upper layer neurons (SATB2+) emerged only at week 12 in the CX-TH assembloids (Fig 2A). The emergence of increased upper layer neurons did not occur at the expense of gliogenesis, as GFAP+ astrocyte population fractions remained similar between conditions (Fig 2B). At weeks 6 and 8, there were significant increases in the fraction of KI67+ dividing progenitors in CX-TH assembloids, trending back to CX-CX levels by week 12 (Fig 2C). In rodents, TCAs have been shown to promote radial glia proliferation through secreted signaling (*9, 10, 59–62*). This influence on radial glia division has been linked to a critical period (*60*), matching our observations of an early influence of human TCAs on increased radial glia proliferation. This provides confidence in the utility of this assembloid model to discover human features of TCA impacts upon cortical cell fate.

**Figure 2.**
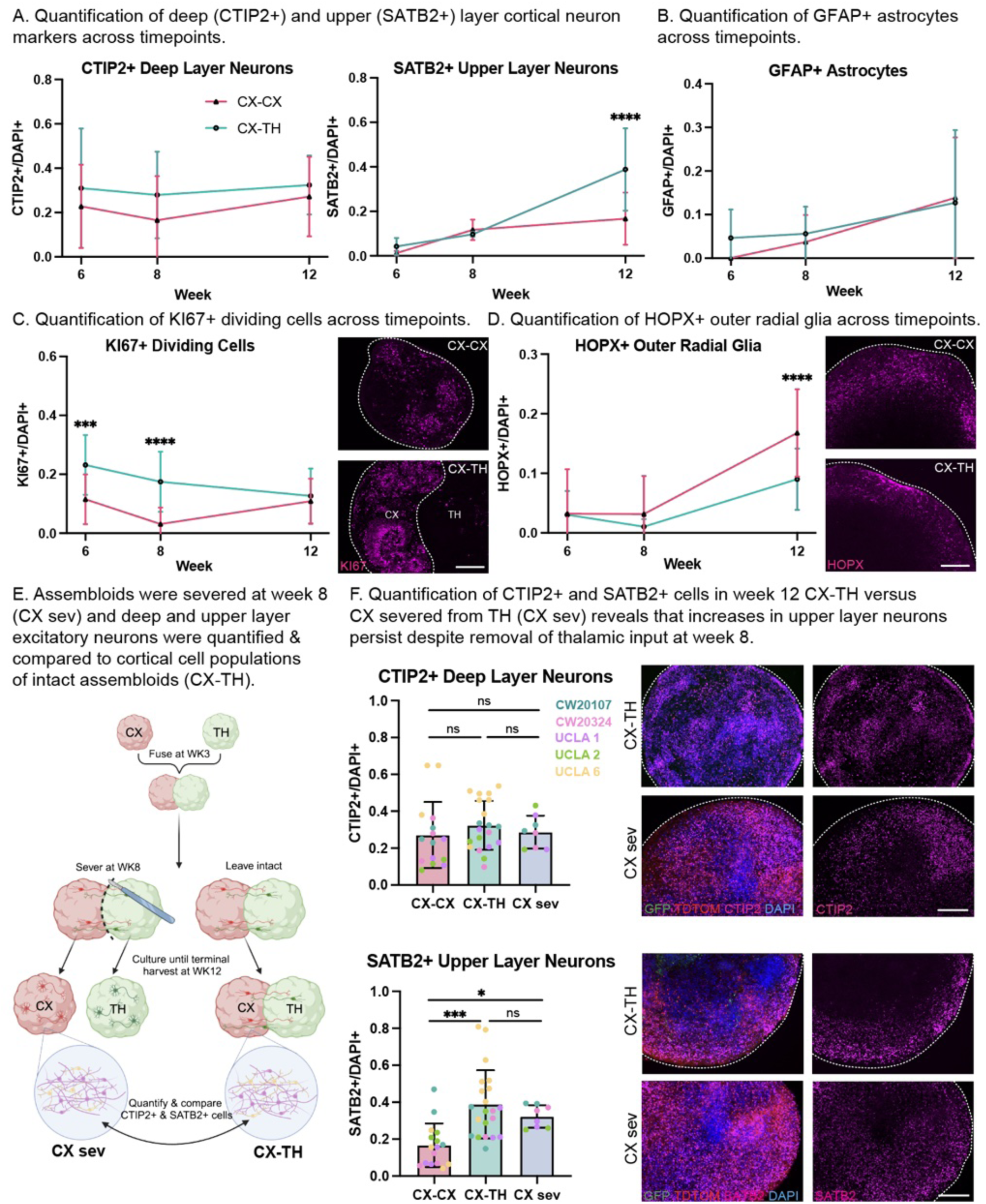
Quantification of cell identity markers across timepoints and assembloid severance suggest that thalamic input in early neurogenesis is necessary and sufficient to drive upper layer specification. A) Quantification of deep (CTIP2+) and upper layer (SATB2+) neurons in TC assembloids reveals a significant difference between CX-TH and CX-CX SATB2+ neurons at week 12 (two-way ANOVA, Tukey multiple comparisons; adj p < 0.0001) but not weeks 6 (adj p = 0.4252) or 8 (adj p = 0.6362). However, no statistically significant differences in CTIP2+ deep layer neurons were observed between assembloid conditions at any time points (weeks 6, 8, 12 adj p = 0.2470, 0.1309, 0.4414, respectively). Cell proportions (y-axis) are expressed as the total number of cell identity marker+ cells over total DAPI+ nuclei. B) GFAP-positivity was also quantified as a ratio over the total number of DAPI+ cells at weeks 6, 8, and 12. Two-way ANOVA followed by Tukey multiple comparisons demonstrates no statistically significant differences in the relative abundance of astrocytes between assembloid conditions (weeks 6, 8, 12 adj p = 0.2318, 0.6155, 0.7637, respectively), suggesting that thalamic input does not impact the onset of gliogenesis. C) Dividing cells were quantified across time as a ratio of KI67+ cells over total DAPI+ nuclei. Two-way ANOVA followed by Tukey multiple comparisons test revealed that in the CX-TH condition, there were significantly more dividing cells at weeks 6 (adj p = 0.0009) and 8 (adj p < 0.0001) than in the CX-CX condition (week 12 adj p = 0.5923). This suggests that thalamic input may drive changes in cell fate specification by impacting progenitor division. Representative KI67 staining from week 8 UCLA 6 CX-CX (top) and CX-TH (bottom) assembloids are shown (scale bar = 250 μm). In CX-TH, the cortical half of the assembloid, used for quantification, is delineated with a dotted line. D) HOPX+ oRG were quantified across time as a ratio over total DAPI+ nuclei. Analyses reveal that at week 12, CX-CX has significantly more HOPX+ oRGs compared to CX-TH (two-way ANOVA, Tukey multiple comparisons, adj p < 0.0001). No statistically significant differences were observed at weeks 6 (adj p = 0.9240) or 8 (adj p = 0.3051). This indicates that thalamic input may be pushing cortical oRGs to differentiate into upper layer neurons such that there are fewer of these cells at the week 12 timepoint in CX-TH conditions. E) Schematic of TC assembloid severance protocol. Thalamic and cortical organoids were fused at week 3, then TC assembloids were severed with a scalpel at week 8, after which cortical and thalamic halves were cultured separately in 50/50 media. At week 12, both severed and intact assembloids were fixed and cortical halves stained for canonical deep (CTIP2+) and upper (SATB2+) layer cortical neurons. Generated using BioRender. F) CTIP2+ and SATB2+ cells in cortical organoids, in severed (CX sev) or fused (CX-TH) cortical organoids, were quantified to determine the fraction of deep and upper layer neurons, respectively. Results, with significance calculated by one-way ANOVA followed by Tukey multiple comparisons, demonstrate that severance of the assembloid at week 8 did not significantly change the fraction of deep (adj p = 0.8202) or upper layer neurons (adj p = 0.5397) compared to the CX-TH condition, indicating that the effects of thalamic signaling on cortical upper layer neuron specification were able to persist despite severance. The fraction of SATB2+ neurons in CX sev was significantly greater than that of CX-CX (adj p = 0.0486), while CTIP2+ neurons remained unchanged between these conditions (adj p = 0.9659). Left: Bar graphs depicting the proportion of CTIP2+ or SATB2+ cells across 5 cell lines, normalized to total DAPI+ cells. CX sev data is compared to CX-TH and CX-CX data, repeated from Figure 1C, as severance experiment was performed in parallel. Technical replicates of 2 organoids from each of 4 lines were quantified to achieve per condition averages. Right: Representative images of week 12 cortical organoids in the CX-TH or CX sev conditions. Images in first column show merged channels for GFP+ thalamic afferents (green), tdTomato+ cortical cells, DAPI nuclear stain (blue), and either CTIP2+ deep layer cortical neurons (top panel) or SATB2+ upper layer cortical neurons (bottom panel). Images in second column isolate CTIP2 and SATB2 channels, respectively. Scale bar = 250 μm.

We next explored whether outer radial glia (oRG), a specialized population of progenitors, expanded in humans and nonhuman primates compared to rodents (*14, 17*), were altered with thalamic input. In the CX-TH assembloid, there was a 1.9-fold decrease in HOPX+ oRG at week 12, but not weeks 6 or 8 (Fig 2D). This is corroborated by a decrease in week 12 radial glia at the transcript level (Fig 1D). Furthermore, the average increase in upper layer neurons is reflected in a proportional, reciprocal decrease in HOPX+ oRG, suggesting thalamic input promotes these progenitors to give rise to SATB2+ upper layer cells earlier than in CX-CX assembloids.

To test this hypothesis, we leveraged the *in vitro* nature of our experimental model to explore whether early communication between TCAs and radial glia can permanently alter their cell fate, or whether continuous input from TCAs is required to drive this phenotype. Severance of TCAs to the cortex has been performed *in vivo* in rodents, though *Gbx2* null mice without these afferents are lethal postnatally (*63*). Although induced models of severance result in altered area boundaries and layer 4 maturation (*7, 8, 10, 64–66*), strong links between thalamic and cortical calcium wave patterns make segregation of these regional network maps difficult to decipher (*6*). Thus, true severance with limited cortical off-target effects is challenging *in vivo*, but is fully tractable in the assembloid system.

Thus, we designed an experiment in which TC assembloids were severed at week 8 and cortex and thalamus were grown independently until week 12 (Fig 2E, Fig S4). This severance timepoint, prior to the rise of SATB2+ populations in the CX-TH condition, enables us to test whether transient TCA input is sufficient to drive the upper layer neurogenesis phenotype or whether sustained communication is necessary. At week 12, immunostaining of cortex severed from thalamus (CX sev) was compared to intact CX-TH cortical regions to determine if the increase in SATB2+ upper layer neurons was preserved (Fig 2E). Quantification of both CTIP2 and SATB2 revealed that the increase in upper layer neurons was not affected by severance, while deep layer neurons remained unchanged as in the unsevered condition (Fig 2F). These data suggest that the impact on upper layer neurogenesis mediated by TCA to cortex interaction occurs early; this motivated us to further investigate potential TCA - radial glia communication.

### Physical interactions exist between human thalamic projections and outer radial glia

We next sought to investigate the nature of the TCA - radial glia interaction in greater detail. First, we set out to identify the cell types that TCAs contact during human cortical development. In the assembloid, we observed that thalamic afferents exhibited targeted pathfinding around cortical organoid rosettes that model the ventricular zone, recapitulating previous findings (*34*). However, we also observed that at weeks 6 and 12 of differentiation, TCAs were found intermingled with progenitor cells positive for HOPX, a canonical marker of oRG (Fig 3A, Fig S5). We corroborated this observation in primary developing human cortical tissue at gestational week 20 where we observed canonical GBX2+ TCAs (*64, 67*) in the outer subventricular zone (OSVZ), a cortical lamina rich with oRG (Fig 3B). TCAs were also observed in expected regions of contact, including the intermediate zone and cortical plate (Fig 3B) (*1–3, 19, 24, 68*). These observations highlight an evolutionary divergence in human cortical development that is difficult to probe in the rodent system. Three-dimensional confocal image reconstruction of the assembloid and human developing cortex showed direct, physical contact between the TCAs and HOPX+ cortical oRG in both *in vitro* and *in vivo* contexts (Fig 3C - D).

**Figure 3.**
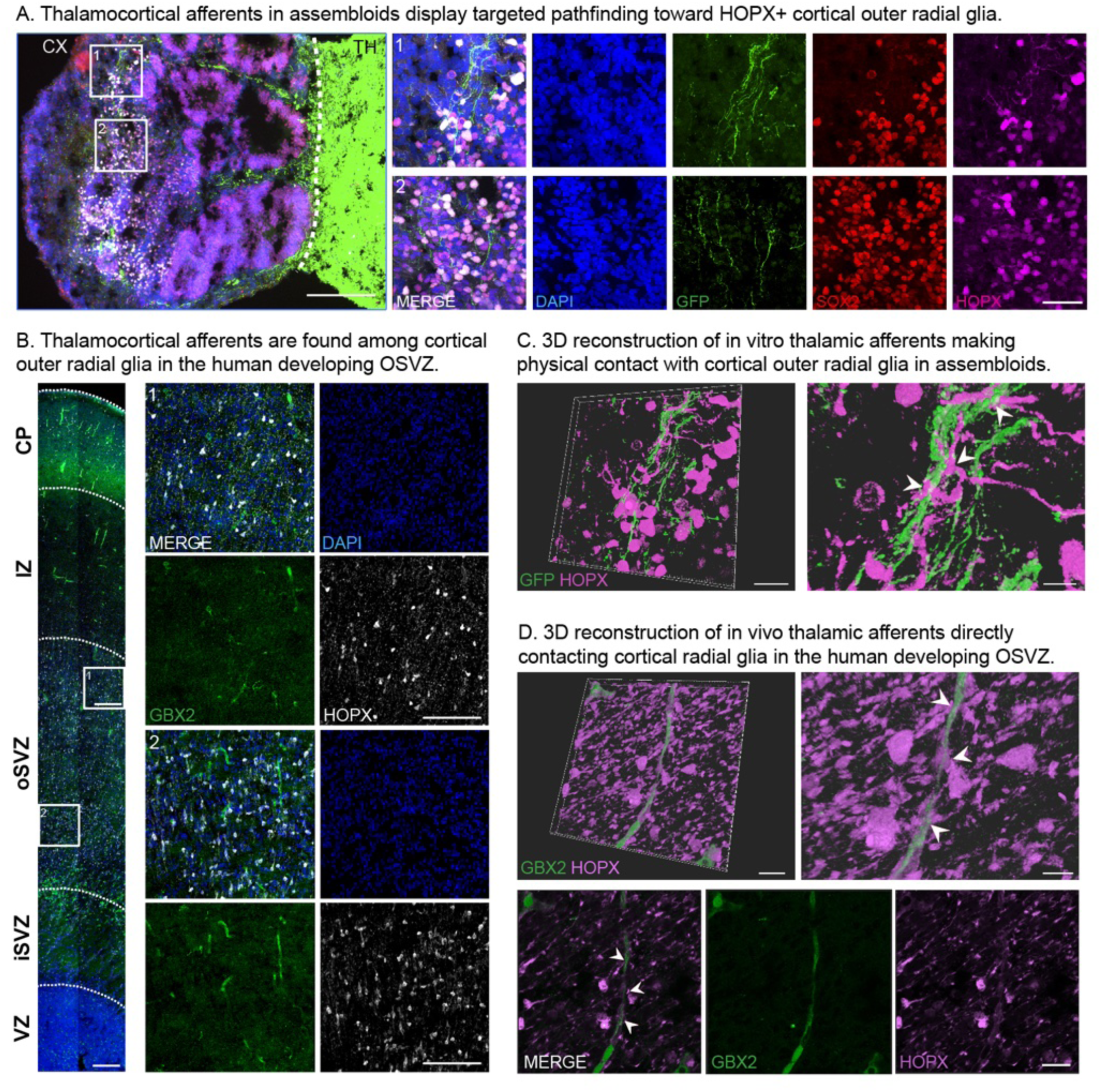
Thalamic afferent tracing demonstrates a physical interaction between thalamic projections and cortical radial glial progenitors in vitro and in vivo. A) Tracing of reciprocal afferents in immunostained TC assembloids reveal that thalamic afferents (GFP+) exhibit targeted pathfinding around cortical organoid rosettes, circular structures that mimic the *in vivo* ventricular zone and contain ventricular radial glia marked by red SOX2 (left, scale bar = 250 μm). GFP+ TCAs were frequently found among HOPX+ cortical oRG, which are typically located in the area surrounding rosettes at week 6. Insets 1 & 2 (right) highlight areas where GFP+ thalamic afferents are found in cortical regions dense with magenta HOPX+ oRG (scale bar = 50 μm). Staining for SOX2, which marks cortical progenitors broadly, demonstrates that TCAs do not appear to target ventricular progenitors to the same extent as they do oRG. Merged images depict cells double positive for SOX2 and HOPX as white. Representative images are from a week 6 TC assembloid generated from the UCLA 6 stem cell line. B) Confocal imaging of gestational week 20 primary human cortical tissue reveals that TCAs are found among oRG of the OSVZ. Left: Tilescan of the span of the cortex with individual layers delineated with dotted lines and the following abbreviations: cortical plate (CP), intermediate zone (IZ), outer subventricular zone (OSVZ), inner subventricular zone (iSVZ), ventricular zone (VZ). GBX2+ thalamic afferents are shown in green, while HOPX+ oRG are depicted in white. Scale bar = 100 μm. DAPI+ nuclei are colored blue. Right: Insets from the OSVZ of the left tilescan, highlighting regions where GBX2+ thalamic afferents are found among HOPX+ oRG (scale bar = 50 μm). C) 3D reconstruction of immunofluorescence staining from Figure 3A, inset 1, depicting GFP+ thalamic axons (green) physically contacting cortical HOPX+ oRG (magenta); left scale bar = 20 μm. Right depicts a zoomed in view with white arrows pointing to areas where thalamic afferents and oRG projections exhibit physical contacts (scale bar = 5 μm). D) 3D reconstruction of immunostaining in gestational week 20 human cortical OSVZ, with purple HOPX+ oRG found making physical contacts with green GBX2+ thalamic axons (scale bar = 20 μm). Right depicts a zoomed in view with white arrows highlighting areas where oRG afferents are found making direct contact with the thalamic axon (scale bar = 8 μm). Bottom row depicts the original confocal maximum intensity projection from which the reconstruction was generated, with channels separated and arrows pointing to the same TCA - oRG contact points (scale bar = 20 μm).

In contrast to lissencephalic rodent models, ferrets have gyrencephalic brains with ample oRG (*69*). Previous studies have shown that binocular enucleation in ferrets disrupts TCAs (*70*), and this model has been used to demonstrate that impeding TCA development inhibits intermediate progenitor cell proliferation and cortical surface area expansion (*71*). This alludes to an evolutionary divergence in which cortical expansion and gyrencephaly may be associated with a distinct mode of communication between TCAs and cortical progenitors. This is consistent with studies showing that TCAs enter the human and nonhuman primate cortex earlier than in rodents and through progenitor zones, though prior anatomical studies were performed early in the second trimester before oRG emerge (*16, 22*). Our findings suggest that this early entry of TCAs into the human cortex enables physical contact with oRG, and we sought to explore the mechanism of this interaction and its functional significance.

### NRXN1 and NLGN1 colocalize with TCA – outer radial glia populations

We next sought to explore the interaction between TCAs and radial glia at the molecular level. Differential gene expression analysis of snRNA-seq data showed that thalamic input in CX-TH assembloids impacts transcriptional programs across all cell types as compared to CX-CX (Fig S6A). Changes in mature neuronal populations were observed and expected; deep and upper layer excitatory neurons from CX-TH had greater upregulation of genes associated with synapse and nervous system development gene ontology (GO) terms at week 6 (Fig S6C). By week 12, gene expression differences between CX-TH and CX-CX became more significant, and GO terms highlight upregulation of transcriptional, translational, and metabolic programs in the cortex with the addition of thalamic input (Fig S6C). Interestingly, we also observe a slight shift in increased prefrontal cortex areal identity in mature excitatory neurons from CX-TH compared to CX-CX (Fig S6B). Most surprisingly, cortical progenitors, such as radial glia, also demonstrate numerous differentially expressed genes. The radial glia cluster had significant upregulation of translational programs at week 6 and programs related to neuronal maturation and neurite outgrowth at week 12 (Fig 4A). This transcriptional data is consistent with our observations that early TCA input modulates radial glia cell fate (Fig 2), and paired with our observation of a physical interaction between TCAs and oRG (Fig 3), we searched for protein candidates that mediate this intercellular contact.

**Figure 4.**
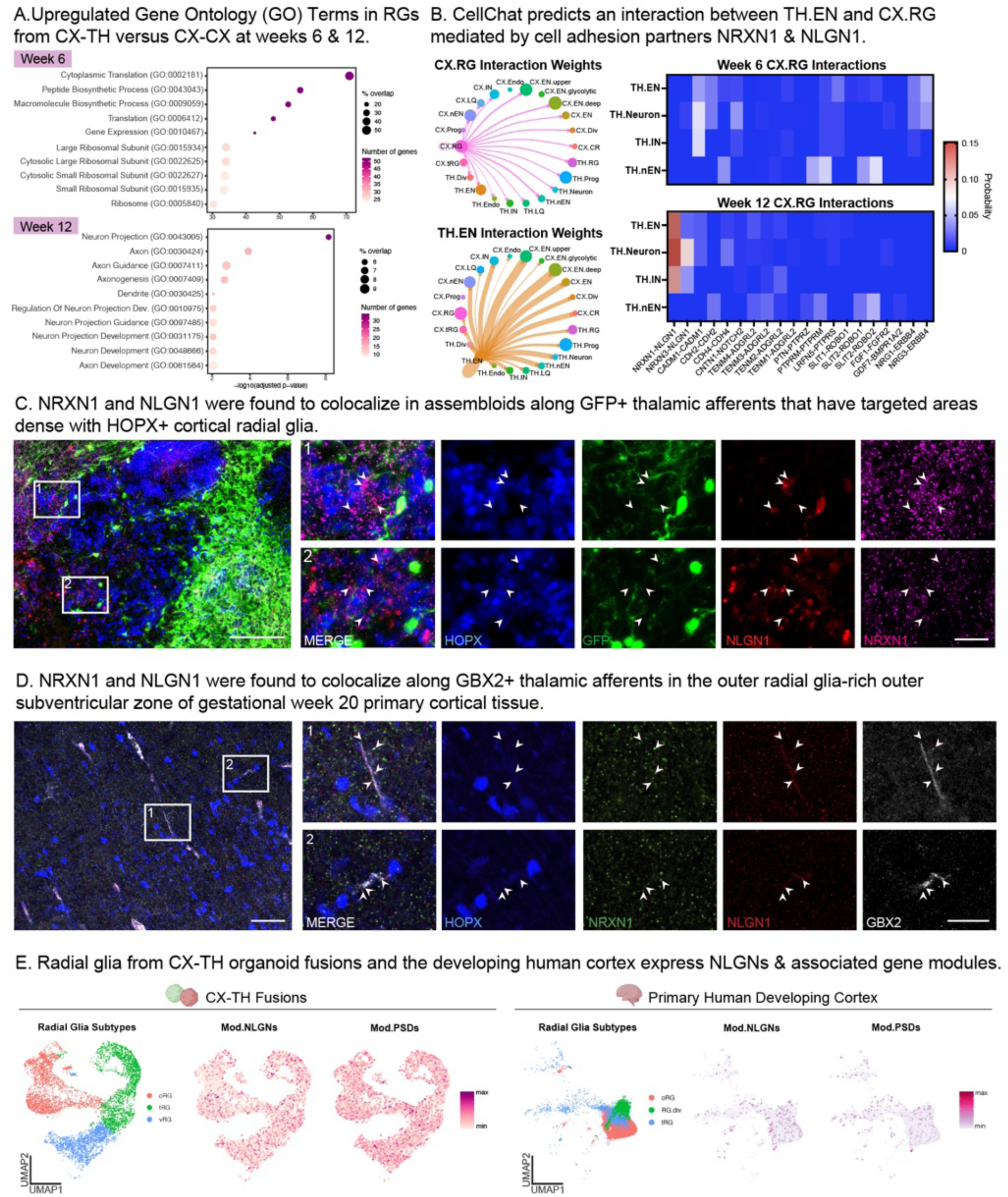
Ligand-receptor analysis identifies the NRXN1-NLGN1 pair as a potential mediator of the thalamic afferent – cortical radial glia interaction, which was validated in organoid and primary tissue. A) Genes upregulated in cortical radial glia in CX-TH compared to CX-CX at weeks 6 and 12 were assessed for gene ontology (GO) term enrichment. At week 6, radial glia (RG) that had access to thalamic input showed greater expression of genes associated with cytoplasmic and ribosomal translation and biosynthetic processes. At week 12, gene expression in this RG population shifted towards GO terms associated with neuron projection, axonogenesis, and neurite development. Circle location on the x-axis depicts significance, while circle color represents number of genes represented and circle size represents percent overlap with the published GO term gene list. B) Left: Circle plots generated using CellChat depict the ligand-receptor interaction weights between cortical radial glia and all other cell types (top), as well as thalamic excitatory neurons and all other cell types (bottom). These graphs demonstrate that CellChat is able to predict expected cell-cell interactions, such as those between thalamic and cortical excitatory neurons that are well described in the literature, as well as novel interactions that can be further explored. This single-nuclei transcriptomic analysis corroborates the possible receptor-ligand interactions between cortical radial glia progenitors and thalamic excitatory neurons, originally detected by immunofluorescence of CX-TH assembloids (Fig 3A). Right: Heatmaps depict the relative CellChat probability weights of various ligand-receptor pairings (x-axis) between cortical radial glia and thalamic neuron subtypes (y-axis). Heatmaps for interaction pair probabilities are shown for both week 6 (top) and week 12 (bottom). Heatmaps highlight that while interactions between cortical radial glia and thalamic neurons are predicted across timepoints, the NRXN1-NLGN1 pair appears to be the most likely interaction, having row maximum probability scores from 0.10 - 0.15. C) Representative image of the cortical half of a TC assembloid at week 6 from line CW20324, where NRXN1 (magenta) and NLGN1 (red) are found to colocalize along GFP+ (green) thalamic afferents that are contacting HOPX+ (blue) cortical oRG (scale bar = 250 μm). Insets depict zoomed in views of these colocalizations, with arrows pointing to areas where NRXN1 and NLGN1 puncta overlap (scale bar = 50 μm). D) *In vivo* NRXN1 and NLGN1 colocalization was validated in the OSVZ of gestational week 20 primary cortical tissue. Image at right depicts cortical OSVZ primarily populated with HOPX+ (blue) oRG in GW20 primary tissue. In this image, NRXN1 (green) and NLGN1 (red) are found to colocalize along GBX2+ (white) thalamic afferents that are contacting HOPX+ cortical oRG (scale bar = 250 μm). Insets 1 & 2 highlight colocalizations, with arrows pointing to areas where NRXN1 and NLGN1 puncta overlap (scale bar = 50 μm). E) Feature plots showing expression of neuroligins (Mod.NLGNs) and related scaffolding proteins (Mod.PSD) in radial glia cells from CX-TH assembloids and developing primary human cortex.

Using our snRNA-seq data across 5 cell lines, we performed a receptor-ligand interaction analysis (*72*) to identify candidate signaling molecules regulating interactions between these cells (Fig 4B). At week 6, few receptor-ligand pairings appeared to be statistically significant, but by week 12, the NRXN1 - NLGN1 pair was predicted to be a likely mediator of interaction between thalamic excitatory projection neurons and cortical radial glia (Fig 4B). NRXN1 - NLGN1 have been extensively described to have roles in the development and stabilization of synapses and tri-partite synaptic interactions (*41–48*), but have also been described to promote cell adhesion and dendrogenesis in non-synaptic contexts (*49, 50, 52*). Our experimental data suggest an early interaction between TCAs and radial glia is essential, yet this computational analysis only identified predicted physical interactions at later time points. We thus sought to validate if NRXN1 and NLGN1 could be colocalized in our organoids, and at what developmental timepoints.

NRXN1 and NLGN1 proteins colocalized adjacent to HOPX+ cortical radial glia and thalamic afferents in the organoid as early as week 6 (Fig 4C) and was maintained through week 12 (Fig S7). This indicates an early and sustained interaction between TCAs and cortical radial glia. We further validated NRXN1/NLGN1 colocalization in primary human cortical tissue from mid-gestation (Fig 4D). We observed colocalization of NRXN1 and NLGN1 proteins along *in vivo* GBX2+ thalamic afferents in the OSVZ, a distinct cortical lamina unique to gyrencephalic mammals and primarily populated by oRG (Fig 4D). To interrogate the transcripts that correspond to this protein expression, we subclustered the radial glia from our assembloid snRNA-seq and from primary developing cortex scRNA-seq (*73*). In both datasets, we observed expected subtypes of radial glia, including oRG, and saw expression of modules related to NLGN and related intercellular contact gene programs (Fig 4E). This analysis further established the prevalence of NLGN expression in cortical oRG across primary human datasets, motivating us to explore whether transcellular communication exists between these cell types.

### Transcellular tracing validates physical communication between TCAs and cortical outer radial glia

We performed anterograde viral tracing to validate a physical connection between the two cell populations (*74*). Prior to fusion, we separately introduced an AAV1 expressing Cre recombinase in the thalamic organoids and a floxed-Myc-BFP-mCherry lentivirus in cortical progenitors, via the radial glia-specific enhancer SOX2 regulatory region 2 (SRR2) (Fig 5A, Fig S8A - B). This allowed us to track which cortical progenitors received inputs from thalamic neurons via Cre-dependent BFP-to-mCherry conversion (Fig 5A). Using this method, we compared cortical organoids fused with either Cre-expressing (“Transcellular”) or uninfected (“Control”) thalamic organoids, and observed mCherry+ cells in the transcellular condition, which overlapped with cortical progenitor markers for outer radial glia (Fig 5B - C). Colocalization of mCherry+ cells with SOX2, which marks neural progenitors broadly, and HOPX, which marks oRG, demonstrate that thalamic neurons contact multiple radial glia subtypes, including oRG.

**Figure 5.**
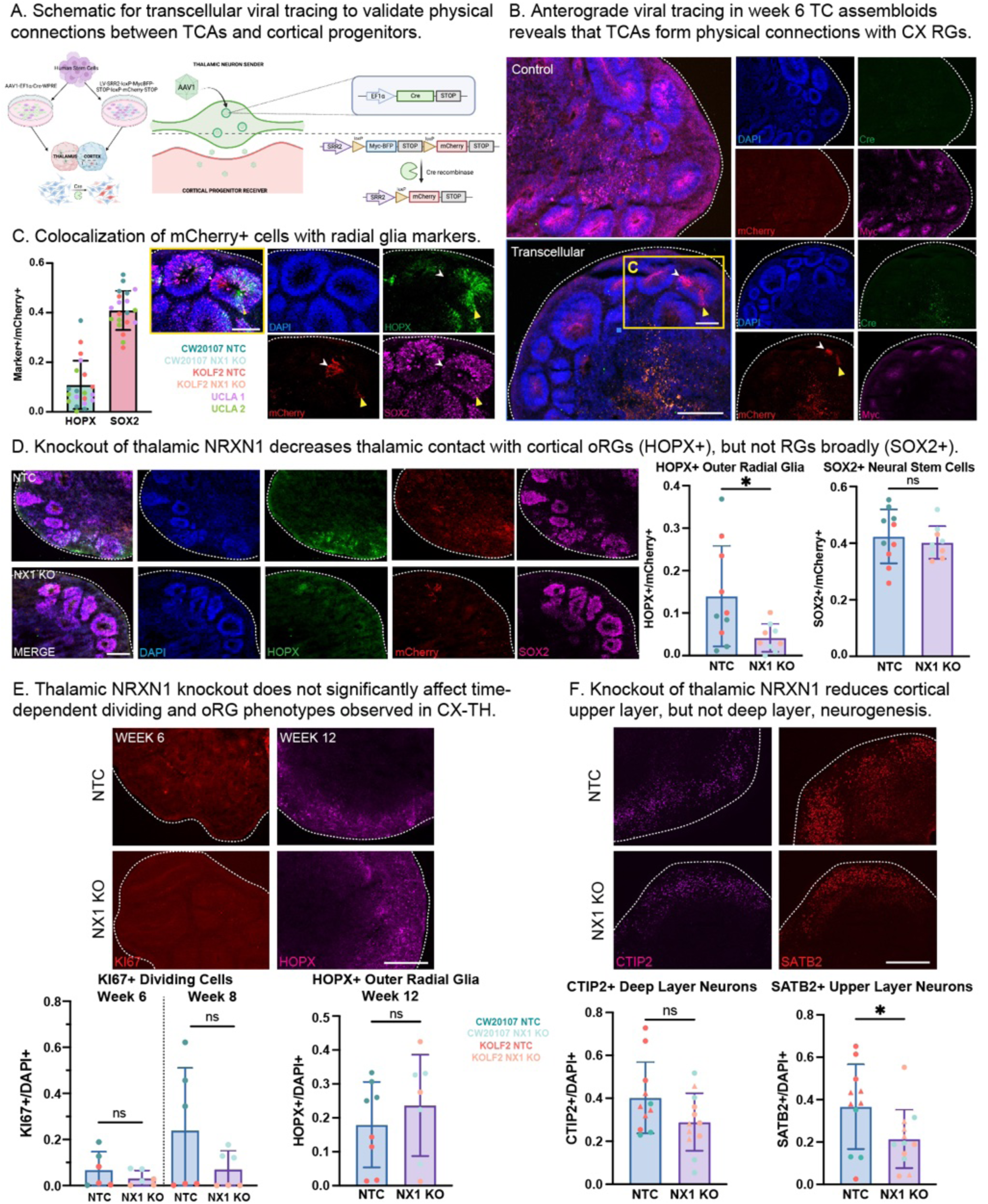
Anterograde viral tracing and knockout of thalamic NRXN1 in thalamocortical assembloids suggest that thalamic contacts with cortical outer radial glia partially mediate upper layer neurogenesis. A) Schematic depicting transcellular AAV1 mechanism of action in TC assembloids, in which mCherry-positivity marks SOX2-expressing (via SRR2 enhancer) cortical progenitors that have received Cre recombinase from thalamic neuronal projections, facilitating a Cre-dependent Myc-BFP to mCherry conversion in fluorescence expression. Generated using BioRender. B) Anterograde viral tracing in week 6 TC assembloids reveals that thalamic axons form physical connections with cortical radial glia. In the “Transcellular” condition, thalamic organoids infected with AAV1-EF1α-WPRE-Cre (green) were fused with cortical organoids infected with a progenitor-specific viral construct LV-SRR2-loxP-Myc-BFP-STOP-loxP-mCherry-STOP, such that when thalamic neurons make contact with cortical progenitors, expression switches from Myc-BFP (Myc, magenta) to Cre-dependent mCherry (red). Arrows indicate regions with mCherry+ cells, with the yellow inset (“C”) further investigated for colocalization (Fig 5C). This is contrasted with the “Control” condition in which an uninfected thalamic organoid is fused with a LV-SRR2-infected cortical organoid to control for any spurious conversion of the lentivirus to mCherry expression due to recombination. Scale bar = 250 μm. C) Quantification and 20X image depicting transcellular mCherry+ (red) cells that colocalize with primarily SOX2 (magenta), with a subset also being positive for HOPX (green). This demonstrates that thalamic neurons are capable of making physical contacts with cortical progenitors, as the transfer of Cre recombinase facilitates the subsequent transition from Myc-BFP to mCherry-positivity (scale bar = 125 μm). The mCherry+ cells were colocalized with HOPX (oRG) and SOX2 (RG), which revealed that an average of 40.9% of mCherry+ cells overlap with SOX2, while only 10.9% overlap with HOPX. D) Representative images of transcellular TC assembloids generated with non-targeting control (NTC) cortex fused with either NTC or NRXN1 knockout (NX1 KO) thalamus (scale bar = 250 μm). Images depict both merged and individual channels with DAPI+ nuclei (blue), HOPX+ oRG (green), transcellular mCherry (red), and SOX2+ radial glia (magenta). To determine whether thalamic NRXN1 knockout impacted the cortical cell types that thalamic axons contacted, overlap of HOPX+ and SOX2+ with mCherry+ was quantified across NTC and NX1 KO conditions. NRXN1 knockout significantly decreased the number of mCherry+/HOPX+ cells (t-test, p = 0.0281), but not colocalization of mCherry with SOX2 (t-test, p = 0.5579). This demonstrates that thalamic NRXN1 knockout decreased the percentage of contacts that thalamic neurons made with cortical oRG, but not radial glial progenitors broadly. E) Thalamic NRXN1 knockout does not appear to significantly affect time-dependent dividing and oRG phenotypes observed in CX-TH compared to CX-CX. Quantification of immunofluorescence cell identity markers demonstrate a non-significant decrease in KI67+ dividing cells at weeks 6 (t-test, p = 0.3143) and 8 (t-test, p = 0.1748). Additionally, no significant increase in cortical HOPX+ oRG was detected in the NX1 KO condition as compared to NTC (t-test, p = 0.4365). F) Cortical deep and upper layer neurons were quantified at week 12 following fusion with either non-targeting control (NTC) or NRXN1 knockout (NX1 KO) thalamic organoids. Dots depict the ratio of marker+ cells over total DAPI+ cells for each condition. Results demonstrate that while no statistically significant difference is observed in CTIP2+ deep layer neurons (t-test, p = 0.0853), thalamic NRXN1 knockout does decrease the average number of cortical SATB2+ cells (t-test, p = 0.440). This suggests that thalamic NRXN1 is important in the promotion of cortical upper layer neurogenesis in TC assembloids (scale bar = 250 μm).

### NRXN1 communication with outer radial glia mediates upper layer neurogenesis

We next sought to determine whether the interaction between NRXN1 and NLGN1 regulates communication between TCAs and cortical radial glia. To do so, we knocked out NRXN1 from 2 stem cell lines and 3 clones (Fig S8E - F) and generated assembloids between NRXN1 knockout (NX1 KO) or non-targeting control (NTC) thalamic organoids and NTC cortical organoids. We observed that in the absence of thalamic NRXN1, there was a 3.5-fold decrease in the transcellular tracing in HOPX+ oRG, but not in radial glia broadly as marked by SOX2 (Fig 5D). This is consistent with the physical interaction that we observed between TCAs and oRG (Fig 3), and suggests that NRXN1 mediates a cell-type specific interaction between TCAs cortical outer radial glia. However, NX1 KO thalamic organoids did not significantly affect the number of dividing cells in their cortical assembloid partners at weeks 6 & 8, nor in the fraction of HOPX+ oRG at week 12 compared to assembloids generated with NTC thalamic organoids (Fig 5E). This data suggests that NRXN1 mediates thalamic contacts with oRG, but does not explain aspects of radial glia division and outer radial prevalence observed in our TC assembloids (Fig 2C - D). Similarly, we saw no significant difference in the fraction of CTIP2+ deep layer neurons in the NX1 KO assembloids. To explore if this interaction is indeed required to mediate the increase in upper layer neurons, we stained for cortical SATB2+ cells at week 12 in assembloids with thalamic NX1 KO or NTC conditions. We saw a 1.7-fold decrease in cortical neurons marked by SATB2 (Fig 5F), leading us to conclude that thalamic NRXN1 is necessary for cortical upper layer neurogenesis via physical interaction with cortical oRG.

## Discussion

In this study, we leverage a 3D *in vitro* model of human thalamocortical development to investigate the role of thalamic afferents in cortical cell fate specification. Our modified human assembloid model provides a tractable system to explore this question as it recapitulates aspects of human brain development and enables the systematic manipulation of thalamic input. We demonstrated that extrinsic thalamic input is necessary and sufficient to drive upper layer neurogenesis, as marked by SATB2-positivity in our assembloid system. When compared to CX-CX assembloids, cortex fused with thalamus (CX-TH) had a significantly greater expansion of upper layer neurons across multiple stem cell lines from varying genetic backgrounds. This evidence builds upon what has previously been noted in mice and rats, which have demonstrated a role for TCAs in promoting expansion of superficial cortical layers, and more specifically layer 4 (*10, 61, 65*). However, unlike findings in traditional rodent models, human cells do not seem to exhibit a tradeoff between upper and deep layer neurons, as expansion of SATB2+ populations did not arise at the expense of deep layer neurogenesis as marked by CTIP2-positivity. Instead, upper layer expansion correlates with decreases in HOPX+ outer radial glia (oRG) at week 12, suggesting that SATB2+ cells may arise from maturation of this progenitor population. This is consistent with observations that humans and non-human primates exhibit cortical expansion in comparison to lissencephalic animals, including mice and rats (*14–16*).

Cortical oRG are also strongly associated with cortical expansion across evolutionary time, and the emergence of the OSVZ comprised almost entirely of these progenitors corresponds with the emergence of gyrencephalic features (*14, 16*), with the largest developmental fractions of these neural stem cells being observed in humans and non-human primates (*14, 17*). The presence of oRG in human stem cell derived cortical organoid models (*57, 75, 76*) provides a unique opportunity to explore TCA input in this evolutionarily distinct context. We demonstrate that an increase in upper layer neurogenesis appears to arise due to early interactions between developing thalamic and cortical cell types. While analysis of cell type temporal dynamics reveals that increases in upper layer neurons do not arise until week 12, severing assembloids at week 8 maintains an increased population of upper layer neurons, despite the removal of thalamic input during mid-cortical neurogenesis. Taken together, these data bolster existing frameworks that implicate upper layer neuronal populations in neurodevelopmental disorders and suggest aberrant development at an early critical period, perhaps in cortical oRG, may confer human vulnerability to these disorders (*77, 78*).

Rodent experiments have shown a clear role for thalamic secreted signals, including Vgf and Wnt3, in driving progenitor division (*9, 10, 59–62*), and our study builds upon these observations by demonstrating specific, physical interactions between TCAs and oRG that mediate expansion of upper layer neurons, another cell type that is expanded in primates compared to rodents (*14, 17*). This physical interaction, which we observe both in organoids and *in vivo* in postmortem developing human cortical tissue, may be enabled by the earlier infiltration of TCAs through progenitor zones of the cortex in primates (*21, 22*), rather than through the intermediate zone to the subplate as seen in mice and rats (*2, 3, 19, 79*). Our observations open additional questions about how other features of these physical interactions may also be functional in modulating human brain development. Although the interactions and phenotypes interrogated here are specific to HOPX+ radial glia, we do observe transcellular communication with SOX2+ radial glia more generally and this may have additional roles in cortical cell fate specification. Previous studies have shown that axons, such as those in the drosophila retina, are capable of driving neurogenesis by acting upon progenitor cells (*80–82*). In rodents, TCA communication with radial glia projections into the intermediate zone have been observed, suggesting our observation of physical contact in the OSVZ is an extension of this existing evolutionary interaction (*59*). Moreover, although the thalamic NRXN1 knockout significantly reduces transcellular communication with HOPX+ oRG, resulting in fewer SATB2+ neurons in the cortex, it is an incomplete reversal of the division and oRG abundance phenotypes that we observe. This suggests additional aspects of thalamocortical communication may be implicated, and our study provides a framework to interrogate these features of human brain development using the assembloid system.

Here, we provide evidence for a non-canonical role of neurexins, whereby thalamic NRXN1 facilitates a physical interaction with cortical progenitors. While neurexins are classically characterized as presynaptic cell-adhesion molecules that interact with postsynaptic neuroligin proteins to promote synapse formation (*83–85*), numerous studies have demonstrated that this family of proteins plays additional roles. Furthermore, NRXN - NLGN pairs, which we identify as possible receptor-ligand partners mediating TCA - oRG interactions, have now been described to play developmental roles outside of synaptic contexts, including functioning in aiding axon pathfinding, dendritic branching, and gliogenesis (*86–89*). This is not entirely surprising as NRXN and NLGN family members are expressed across several glial and non-neural cell types (*53, 90–92*). These expanded behaviors of neurexins, and NRXN1 in particular, prompts future investigation of the specific nature of TCA - oRG communication.

Our observation that TCA interactions with human expanded progenitors (oRG) drive human expanded neuronal layers (upper layers) evokes questions of whether this process may explain evolutionary divergence, but also a vulnerability to neurodevelopmental disorders. Indeed, emerging studies of thalamic anatomy demonstrate a dysregulation in neurodevelopmental disorders (*30–33*) and organoid experiments have successfully explored mechanisms of neurodevelopmental disorders with these assembloid systems (*38, 39*). Both NRXN1 and SATB2, which are central to our observations in typical development here, have been implicated as being mutated in certain neurodevelopmental disorders (*93–105*). Thus, our discovery of a novel, non-canonical role for thalamic NRXN1 in mediating cortical neuron specification via interactions with progenitors provides an opportunity to expand our understanding of both typical development and the etiology of disorders with developmental origins.

## Acknowledgments

We would like to thank the members of the Bhaduri Lab for their insightful advice and comments on the study. We would like to thank L. Zipursky and C. Portera-Cailliau for helpful suggestions and feedback. We would like to thank the Broad Stem Cell Research Center Imaging Core, Charina Julian (UCSF Institute for Human Genetics) for help with running sequencing.

## Author contributions

Conceptualization: AB

Methodology: CVN, AM

Validation: CVN, AM

Formal Analysis: CVN, AM, DCJ

Investigation: CVN, AM, PRN, DCJ, JAS, JM, RW, JMM

Resources: AB, DHG

Data Curation: CVN, AM

Writing - Original Draft: AB, CVN

Writing - Revision & Editing: All authors; Visualization: CVN, AM

Supervision: AB

Project Administration: AB

Funding Acquisition: AB, DHG, additional fellowships in acknowledgments

## Methods

### Cell and Tissue Culture

#### Stem Cells

Human hESCs and iPSCs were authenticated at source (Table S1). Stem cells were frozen in mFreSR (Stem Cell Technologies, Cat. No. 05855) and stored long-term in liquid nitrogen. Cells were thawed at 37°C into mTeSR+ medium (Stem Cell Technologies, Cat. No. 100-0276) supplemented with 1X penicillin-streptomycin (Life Technologies, Cat. No. 15140122) and plated onto 6-well plates coated with growth factor-reduced matrigel (Corning, Cat. No. 354230). To limit cell death upon thawing, stem cells were treated overnight with 10 μM Rock inhibitor Y-27632 (Tocris Cat. No. 1254). Stem cells were maintained in mTeSR+ at 37°C, 5% CO2 with media changed every other day. When cells reached 75-90% confluency, they were passaged by washing with PBS, a 1 minute room temperature incubation with ReLeSR (Stem Cell Technologies, Cat. No. 100-0483), followed by aspirating and a 5 minute dry incubation at 37°C. Stem cells were dislodged by adding mTeSR+ and rinsing the well prior to expanding the colonies into 4-6 wells of a matrigel-coated 6-well plate.

#### Viral Infection

Viral constructs were introduced at the stem cell stage during the second passage post-thaw. To do so, a confluent well of stem cells was passaged as described above and virus was added to the media (Table S2). Media was changed to remove virus 18 hours post-infection.

#### Organoid Generation

Differentiation and organoid culturing were based on previously published protocols (*36, 56*). Stem cells were washed with PBS and dissociated into a single-cell suspension using a 5 minute incubation with accutase (Millipore Sigma, Cat. No. A6964) at 37°C. Cells were then scraped and pelleted by centrifugation at 300 g for 5 minutes. Stem cells were then resuspended in media to induce either cortical or thalamic differentiation at a concentration of 100,000 cells per mL. The cell suspension was then plated at 100 uL, or 10,000 cells, per well in a low-attachment 96-well V-bottom plate (S-Bio, Cat. No. MS-9096VZ). While in culture, organoid media changes were performed every 2 - 3 days.

#### Cortical Differentiation

To drive cortical patterning, stem cells were aggregated and cultured for 18 days in GMEM (Life Technologies, Cat. No. 11710-035) supplemented with 20% Knockout Serum (Life Technologies, Cat. No. 10828-028), 1X NEAA (Life Technologies, Cat. No. 11140050), 1X sodium pyruvate (Life Technologies, Cat. No. 11360070), 1X penicillin-streptomycin, 100 µM ß-mercaptoethanol (Life Technologies, Cat. No. 21985023), 20 µM Y-27632, 5 µM SB431542 (Tocris, Cat. No. 1614), and 3 µM IWR1-e (Cayman Chem, Cat. No. 13659). On day 6, Rock inhibitor Y-27632 was removed from the culture media. At day 18, organoids were transferred to low-attachment 6-well plates (Corning, Cat. No. 07-200-601) and placed on an orbital shaker rotating at 100 rpm. From days 18-35, organoids were switched to DMEM/F12 + Glutamax (Life Technologies, Cat No. 10565-018) supplemented with 1X N2 (Life Technologies, Cat. No. 17502-048), 1X CD Lipid Concentrate (Life Technologies, Cat. No. 11905-031), and 1X penicillin-streptomycin. At day 35, cortical culture media was supplemented with additional 10% heat-inactivated FBS (Life Technologies, Cat. No. 10082147), 5 µg/mL heparin (Millipore Sigma, Cat. No. H3149-10KU), and 0.5% matrigel. From day 70 onward, cortical media was additionally supplemented with 1X B27 (Life Technologies, Cat. No. 17504044) and matrigel concentration was increased to 1%.

#### Thalamic Differentiation

Thalamic differentiation was based on a published protocol (*34, 58*) with some slight modifications to parallelize with cortical differentiation protocols established in the Bhaduri Lab. To induce thalamic patterning, aggregated stem cells were cultured from days 0 - 8 in media containing DMEM/F12 + Glutamax and supplemented with 15% Knockout Serum, 1X NEAA, 100 µM ß-mercaptoethanol, 1X penicillin-streptomycin, 100 nM LDN-193189 (Selleckchem Cat. No. 6053), 100 µM SB431542, and 4 ug/mL insulin (Life Technologies, Cat. No. 12585014). 50 µM Y-27632 Rock inhibitor was also supplemented from days 0 - 4 and FBS from days 0 - 2. On day 9, thalamic organoids were switched to a media containing DMEM/F12 + Glutamax, 0.15% dextrose (Life Technologies, Cat. No. A24940-01), 100 µM ß-mercaptoethanol, 1X N2, 2% B27 without vitamin A (Life Technologies, Cat. No. 12587-010), 1X penicillin-streptomycin, 30 ng/mL BMP-7 (Life Technologies, Cat. No. PHC9544), and 1 µM PD0325901 (Selleckchem, Cat. No. S1036). From day 19 onwards, media was switched to a final differentiation media containing a base of 50% DMEM/F12 + Glutamax and 50% Neurobasal Medium (Life Technologies, Cat. No. 21103049), supplemented with 1X N2, 2% B27 without vitamin A 0.5% NEAA, 1% Glutamax, 0.025% insulin, 50 µM ß-mercaptoethanol, 1X penicillin-streptomycin, 20 ng/mL BDNF (Invitrogen Cat. No. RP-8642, and 200 µM L-ascorbic acid (Sigma-Aldrich, Cat. No. A4403-100MG).

#### Fusion Protocol

Thalamocortical (TC) assembloids were generated at week 3 of cortical and thalamic differentiation in parallel. One cortical plus one thalamic organoid were co-plated in each well of a stationary low-attachment 96-well V-bottom plate for 5 days in a culture media composed of 50% cortical and 50% thalamic media (“50/50 media”), by volume. Media changes were performed every 1-2 days, as needed. After 5 days, each assembloid was transferred to individual wells of a 24-well low-attachment cell culture plate and placed back on the shaker. Assembloids were cultured in 50/50 media, with cortical and thalamic components following media timelines outlined above, until terminal harvest at weeks 6, 8, and 12.

### Immunofluorescence

#### Fixation & Embedding

Organoids were fixed in 4% paraformaldehyde (PFA) for 45 minutes at room temperature, washed with PBS, then placed into 30% sucrose solution overnight at 4°C. Following sucrose incubation, organoids were embedded in a mixture of 50% OCT + 50% sucrose solution and stored at -80°C. Embedded, frozen organoids were sectioned at 16 μm and placed onto treated glass slides. Similarly, primary tissue was obtained and flash frozen in OCT as previously described (*106*). 20 μm cryosections were mounted onto treated glass slides, where they were fixed for 2 minutes with 4% PFA then washed 3 x 10 minutes with PBS.

#### Immunostaining

Slides were treated with boiling citrate-based antigen retrieval buffer (Vector Laboratories, Cat. No. H-3300-250) for 20 minutes, then washed with PBS. Slides were subsequently blocked with blocking buffer (5% donkey serum, 3% bovine serum albumin, 0.1% Triton) for 30 minutes at room temperature before incubation with primary antibodies (Table S3), diluted in blocking buffer, overnight at 4°C. A 3 x 10 minute wash with PBST (0.1% Triton in PBS) was used to remove excess primary antibody solution. Slides were then incubated in the dark with secondary antibodies and DAPI diluted in blocking buffer for 2 hours at room temperature. Excess secondary antibody (Table S3) solution was washed off with PBS prior to mounting glass coverslips with prolong gold (ThermoFisher, Cat. No. P36934).

#### Microscopy

Stained slides were imaged using either the EVOS M5000 or Zeiss LSM880. Acquisition parameters were kept constant among sections with antibody panels being used for quantification purposes. Quantification of cell types and colocalization was performed using Imaris image analysis software, version 10 (Oxford Instruments). Fluorescence intensities were adjusted for display purposes only using FIJI ImageJ software (RRID:SCR_003070).

### Cell and Tissue Culture

#### Sample Capture & Sequencing

Organoids were dried by removing all media then flash frozen and stored at -80°C in preparation for single-nuclei RNA-sequencing. Nuclei were extracted from frozen organoids using the 10X Chromium Nuclei Isolation Kit using a 4-minute lysis time. Nuclei were washed twice using the included buffer, counted using a hemacytometer, then resuspended in buffer to achieve a target capture of 10,000 nuclei. During counting, nuclei were also imaged to ensure healthy morphology and lack of debris. For single-cell samples, organoid tissue was dissociated and processed via FACS as described above. Samples were captured using the 10X Genomics Chromium Single Cell 3’ Reagent Kit v3.1 or v4 and 10X Chromium System v3.1 or the PIPseq T2 3’ Single Cell RNA Kit v4.0. All samples were assessed for quality control purposes at both the cDNA and library stages using the Agilent 4200 TapeStation System. Single-nuclei and single-cell libraries were sequenced using the Illumina NovaSeq 6000 or NovaSeq X platform.

#### Quality Control & Analysis

Reads were aligned to the GRCh38 human reference genome. Cell-by-gene count matrices were generated using the 10X Genomics CellRanger pipelines or Fluent Bio PIPseeker pipeline with default parameters. Single-nuclei data was processed using CellBender (*107*), an R package designed to remove background noise from droplet-based single-cell assays. Doublets were predicted and removed from all samples using the DoubletFinder R package (*108*) using default parameters. Nuclei or cells containing a minimum of 500 genes and containing less than 5% mitochondrial gene content were retained for further analysis. Cortical and thalamic cells were segregated bioinformatically using the MapQuery function to compare cells to primary tissue datasets (*73, 109, 110*) and a pilot organoid batch in which thalamic and cortical halves were dissected prior to sequencing. Following normalization and scaling, Principal Component Analysis (PCA) was conducted using the top 2,000 variable genes using the standard Seurat v5 pipeline (*111*). The number of dimensions was determined using previously described methods (*112*). Differentially expressed genes and projections of the organoid data onto primary tissue datasets (*73, 106, 110*) were used to annotate cell type clusters. Uniform Manifold Approximation and Projection (UMAP) was used to visualize clusters in 2D space using the determined dimensions. Seurat object containing cortical and thalamic nuclei data and associated metadata (Table S4) are included in the supplementary materials.

Differentially expressed genes between comparison datasets with an adjusted p-value < 0.01 were used as input data in Enrichr (*113, 114*), a search tool that enables querying of published, annotated gene datasets. GO Terms (Biological Process, Cellular Component, Molecular Function 2025) were aggregated and the top terms were depicted using their - log10(adjusted p-value) for visualization purposes.

Cell-cell and receptor-ligand interactions were predicted using single-nuclei data input to CellChat (*72*), a tool for quantitatively inferring and analyzing intercellular communication networks from transcriptomic data.

Module activity scoring was performed using gene lists obtained from an atlas of mid-gestational primary human cortex (*4*). Rostrally enriched genes (correlation of > .825) and caudally enriched genes (correlation of > .775) were selected from rostral-caudal bulk gene expression gradient values averaged from 15-21 post-conceptional weeks and across all developing cortical layers (VZ, SVZ, IZ, SP, CP, MZ). Using only these genes, area module scores for prefrontal cortex (PFC, rostral) and primary visual cortex (V1, caudal) were calculated based on average expression level of each gene in cortical excitatory neurons in CX-TH and CX-CX. The score was generated using the sum of all normalized counts per million detected for each module gene, which was subsequently divided by the total number of module genes to reduce bias based on the number of genes in each module.

### Virus Generation

#### Transcellular Viruses

The AAV-EF1⍺-Cre-WPRE plasmid was generated by switching the hSyn promoter of pENN.AAV.hSyn.Cre.WPRE.hGH (Addgene, Cat. No. 105553) with EF1⍺. Briefly, the backbone was cut from the original plasmid with ApaI and EcoRI, and an EF1a promoter sequence flanked by homology arms were cloned into the resulting backbone using the NEBuilder HiFi DNA Assembly Master Mix (New England Biolabs Cat No. E2621). To generate AAV-Cre viruses with the AAV1 serotype, Using a molar ratio of 1:1:1 and Lipofectamine 2000 (Invitrogen, Cat. No. 11668019), HEK293T cells grown in DMEM + GlutaMAX-I (Gibco, Cat No. 10569-010) with 10% Heat-Inactivated Fetal Bovine Serum (Omega Scientific, Cat No. FB-12) were transfected with AAV-EF1⍺-Cre-WPRE, pAAV2/1 (Addgene, Cat. No. 112862) for expressing Rep2/Cap1, and pAdDeltaF6 (Addgene, Cat. No. 112867) for expressing adenovirus E4, E2A, and VA. The next day, the original HEK293T media was replaced with serum-free DMEM to improve virus purification efficiency. Three days post-transfection, AAV1-Cre viruses were harvested using the AAVpro Cell and Supernatant Purification Kit Max (All Serotypes) (Takara Bio, Cat No. 6676) following the manufacturer’s protocol.

To generate the Cre-dependent, dual fluorescent, SOX2 enhancer-driven reporter plasmid LV-SRR2-loxP-Myc-BFP-STOP-loxP-mCherry-STOP, the loxP-BFP-mCherry cassette was first PCR-amplified from a loxP-Myc tag-BFP-STOP-loxP-mCherry-NTR plasmid template (Addgene, Cat. No. 108456) and cloned into pDONR221 using Gateway BP Clonase II (Invitrogen, Cat No. 11789020). The coding sequence was subsequently transferred into the Gateway cassette of the pLEX305-SRR2-WPRE destination vector using Gateway LR Clonase II (Invitrogen Cat No.11791020). To prepare lentiviruses, 12 µg of the plasmid was transfected into HEK293T cells along with 3 µg of pMD2.G and 9 µg of psPAX2. Media was replaced the next day, and the supernatant containing viruses were collected for two consecutive days thereafter. Viral particles were concentrated using Lenti-X Concentrator (Takara Bio, Cat. No. 631232) and resuspended in PBS prior to storage.

#### Short-hairpin RNA Knockdown of NRXN1 and NLGN1

To knockdown the expression of NRXN1 and NLGN1, pLKO.1 and pLKO.3G plasmids were obtained (Addgene, Cat. No. 10878, 14748). pLKO.1 was subsequently modified by removing the puroR sequence and replaced with mCherry via NEBuilder HiFi DNA Assembly reaction (New England Biolabs, Cat. No. E2621L). NRXN1 shRNA sequence (5’-CATCGCCATTGAAGAATCCAA-3’) was cloned into the pLKO.3G plasmid, and NLGN1 shRNA sequence (5’-CCTGCTGACTTTATCCCATTA-3’) was cloned into modified pLKO.1 plasmid with mCherry. Additionally, a scramble sequence (5’-CCTAAGGTTAAGTCGCCCTCG-3’), was cloned into both plasmids as an experimental control. Insertion of shRNA sequences into the pLKO.1 and pLKO.3G backbone were performed according to the manufacturer’s instructions from Addgene. NRXN1 and NLGN1 shRNA sequences were designed using the Broad Institute Genetic Perturbation Platform Web Portal.

Generation of lentivirus carrying these plasmids was performed by transfecting HEK293T cells with the pLKO plasmid alongside both psPAX2 and pMD2.G using Lipofectamine 2000 (Invitrogen, Cat. No. 11668019). The viral media produced from this process was concentrated using Lenti-X Concentrator (Takara Bio, Cat. No. 631232) for a final resuspension of lentivirus.

Knockdown was confirmed using RT-qPCR. RNA was isolated from organoids using RNeasy Plus Mini Kits (Qiagen, Cat. No. 74134) and RNA concentrations were measured using Thermo Scientific NanoDrop One Spectrophotometer (Cat. No. 13-400-525). RNA was converted to cDNA SuperScript™ IV VILO™ Master Mix (Invitrogen, Cat. No. 11756500). RT-qPCR reactions were performed by mixing sample cDNA, gene-specific primers, and Power SYBR™ Green PCR Master Mix (Applied Biosystems, Cat. No. 4368708). Reactions were loaded in 384-well optical plates (Applied Biosystems, Cat. No. 4309849) and analyzed using a QuantStudio™ 5 Real-Time PCR System (ThermoFisher, Cat. No. A28140). Each sample-primer pairing was performed in triplicate and averaged. Genes of interest were normalized to the signal of housekeeping gene GAPDH and ΔΔCT values were calculated for comparison to appropriate controls. Primer pairs were as follows:

NLGN1:

Forward: 5’-CAACAGGGGAACGTCGTTTTC-3’

Reverse: 5’-AGCATGACTTCTGGCAATCTG-3’

NRXN1:

Forward: 5’-GTCCGCCTCGCAAGGAT-3’

Reverse: 5’-ATTTCCATGGCAGCAGCAAG-3’

GAPDH:

Forward: 5’-GGAGCGAGATCCCTCCAAAAT-3’

Reverse: 5’-GGCTGTTGTCATACTTCTCATGG-3’

### Generation of NRXN1 Knockout Stem Cell Lines

To generate clonal homozygous knockout lines for NRXN1, a multi-gRNA ribonucleoprotein (RNP) approach was employed. A pool of three gRNAs targeting an early exon of the gene (GCGUGGUGUUGCGGAACUGG, CGCUGAGCCUGCGACGCUCC, GAGCUGAUUCUGACGCGCGG) was designed and purchased from Edit Co’s GKO Revolution Kit.

For RNP assembly, 6 µl of 33 µM gRNA pool was mixed with 2 µl of 20 µM Alt-R™ S.p. Cas9-RFP V3 (IDT, Cat. No. 10008163) and incubated for 10 minutes at room temperature. The resulting RNPs were then added to 500,000 cells suspended in 20 µl of Lonza’s X solution (Lonza P3 Primary Cell 4D X Kit, Lonza, Cat. No. V4XP-3032). Nucleofection was performed using the Lonza Nucleofector 4D, after which cells were resuspended in mTeSR™ Plus medium (Stem Cell Technologies, Cat. No. 100-0276) supplemented with a 1:10 dilution of CloneR™2 (Stem Cell Technologies, Cat. No. 100-0691).

Approximately 90% of the culture exhibited RFP fluorescence 24 hours post-nucleofection. Cells were then resuspended using Accutase (Invitrogen, Cat. No. 00-4555-56) and plated at a density of 1 cell per well in 96-well plates via limiting dilution. Following 10 days of culture, colonies were expanded, and Sanger sequencing was performed to validate the edits and clonality (Forward Primer: AGGCCAGGGTGAGCTTGA; Reverse Primer: GCAGCGGGCTGGAGTTT; Sanger Sequencing Primer: AGGCCAGGGTGAGCTTGAGC).

For quality control, pluripotency of the cell lines was validated using OCT4 and TRA-160 immunofluorescent staining (Table S3). Low-pass whole-genome sequencing was conducted to ensure genomic stability.

## Supplementary Materials

**Supplementary Figure 1.**
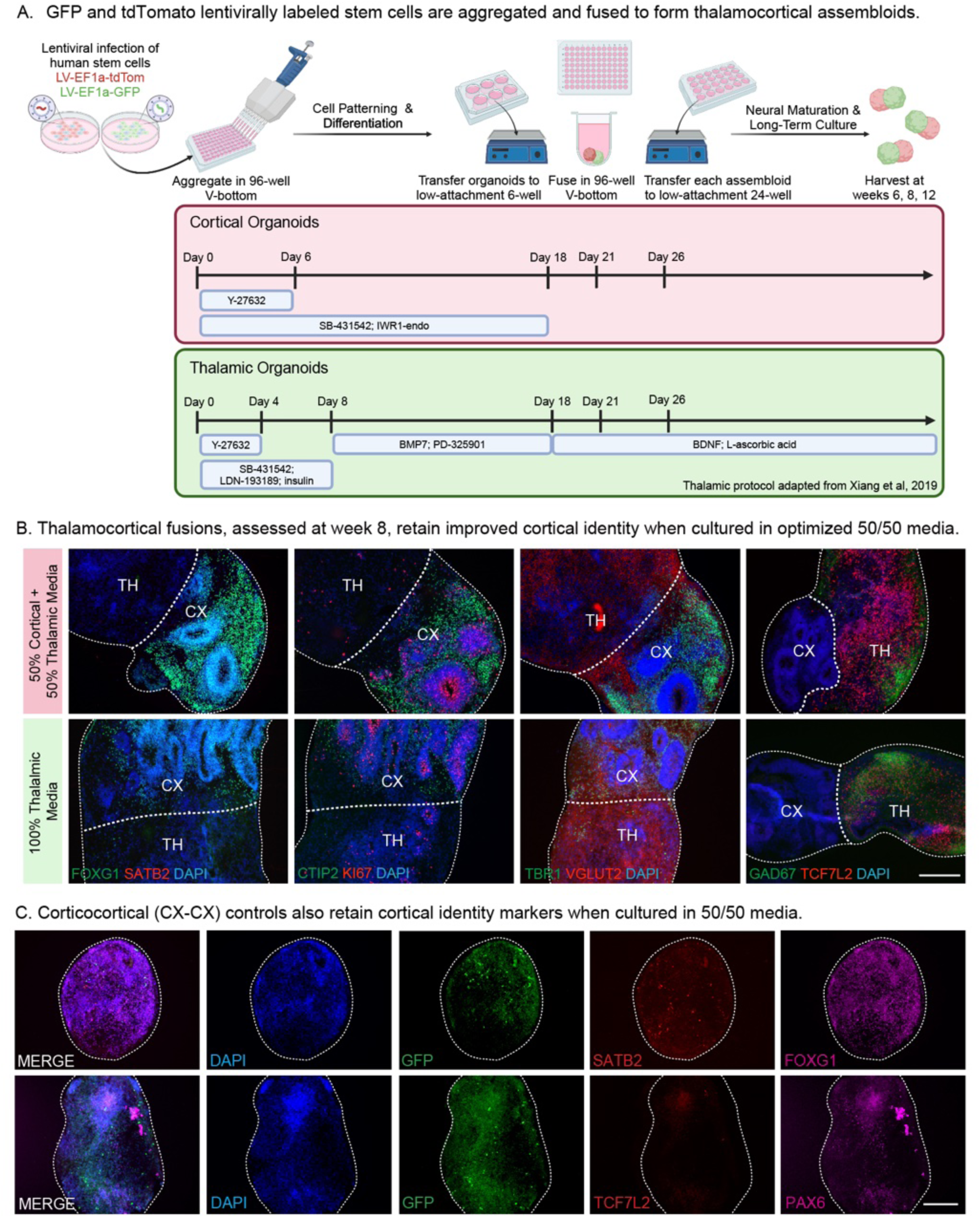
Optimized thalamocortical fusion protocol yields improved cortical identity. A) Workflow of generating TC assembloids. Stem cells are first lentivirally infected with either LV-EF1a-tdTomato (cortical side) or LV-EF1a-GFP (thalamic side) prior to aggregation at day 0. Small molecules are added to the media at designated timepoints to drive cortical and thalamic differentiation. Organoids are transferred to a low-attachment 6-well plate at day 18, then fused at day 21 for 5 days. Assembloids are then transferred to individual wells of a 24-well low-attachment plate and allowed to continue maturing in 50% cortical + 50% thalamic (“50/50”) media until terminal harvest time points at weeks 6, 8, and 12. Generated using BioRender. B) Assembloids cultured in either 50/50 media or only thalamic media were harvested and immunostained for cell identity markers at week 8. Immunofluorescence images demonstrate marked improvement in the conservation of cortical identity when assembloids are grown in 50/50 media, as compared to thalamic media only. 50/50 media conditions demonstrate more robust expression of important cortical markers, such as FOXG1 (progenitor), KI67 (dividing), as well as deep layer cortical markers CTIP2 and TBR1. Importantly, thalamic markers, VGLUT2 (glutamatergic), GAD67 (GABAergic), and TCF7L2 (pan-thalamic) remain enriched in the non-cortical areas of the assembloid. SATB2 (upper layer neurons) are sparsely observed at the week 8 time point in either media condition, as expected. Scale bar = 250 μm. C) Corticocortical (CX-CX) assembloids were used as controls to ensure that any results of the TC assembloids were due to the addition of thalamus, rather than the fusion process itself. CX-CX were cultured in 50/50 media and staining at week 8 showed retainment of important cortical identity markers, such as SATB2 (CX upper layer), FOXG1 (progenitor), and PAX6 (radial glia), but remain negative for thalamic-identity marker TCF7L2. Scale bar = 250 μm.

**Supplementary Figure 1.**
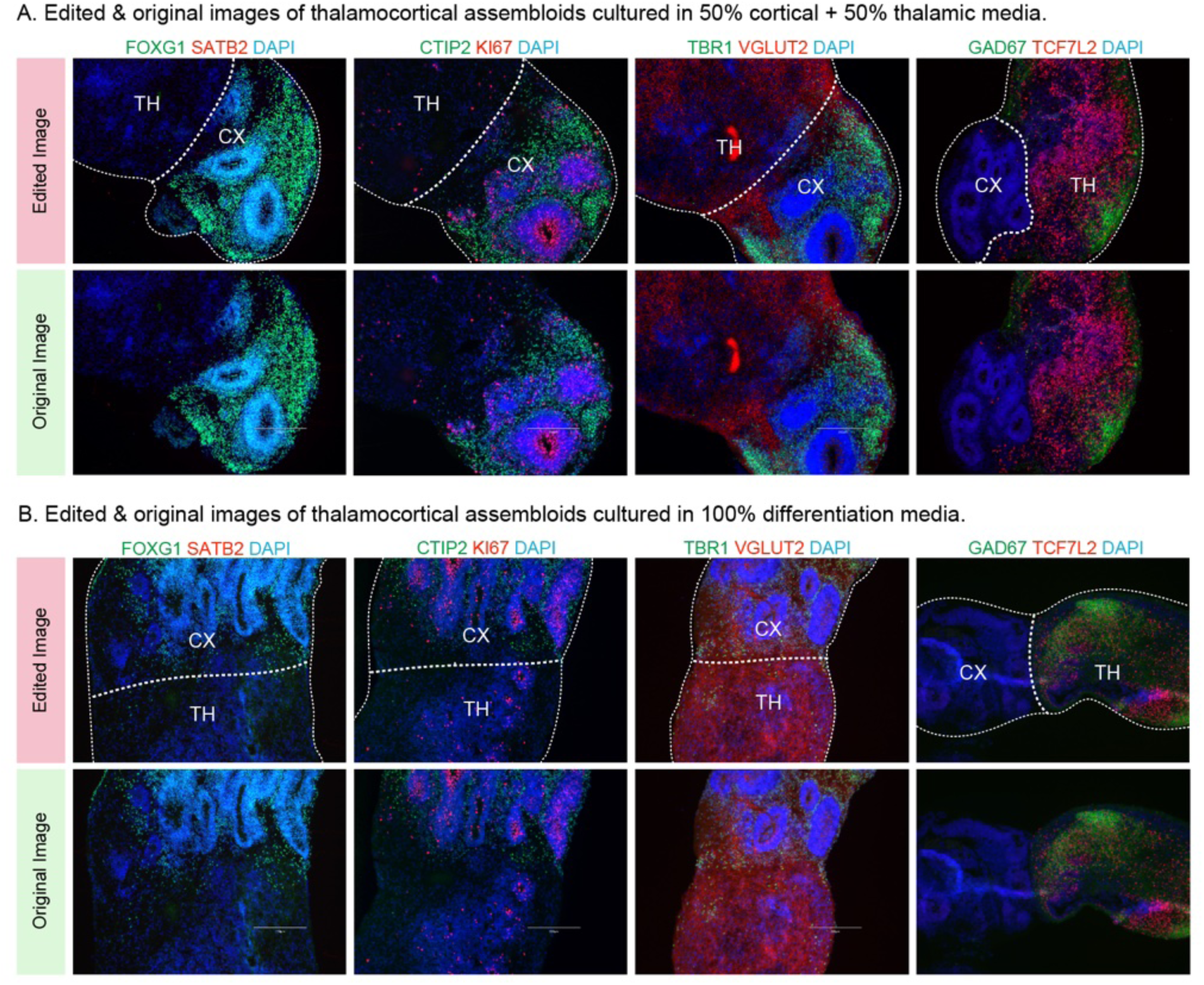
Addendum. A subset of immunofluorescence images from Supplementary Figure 1B were edited to remove scale bars.

**Supplementary Figure 2.**
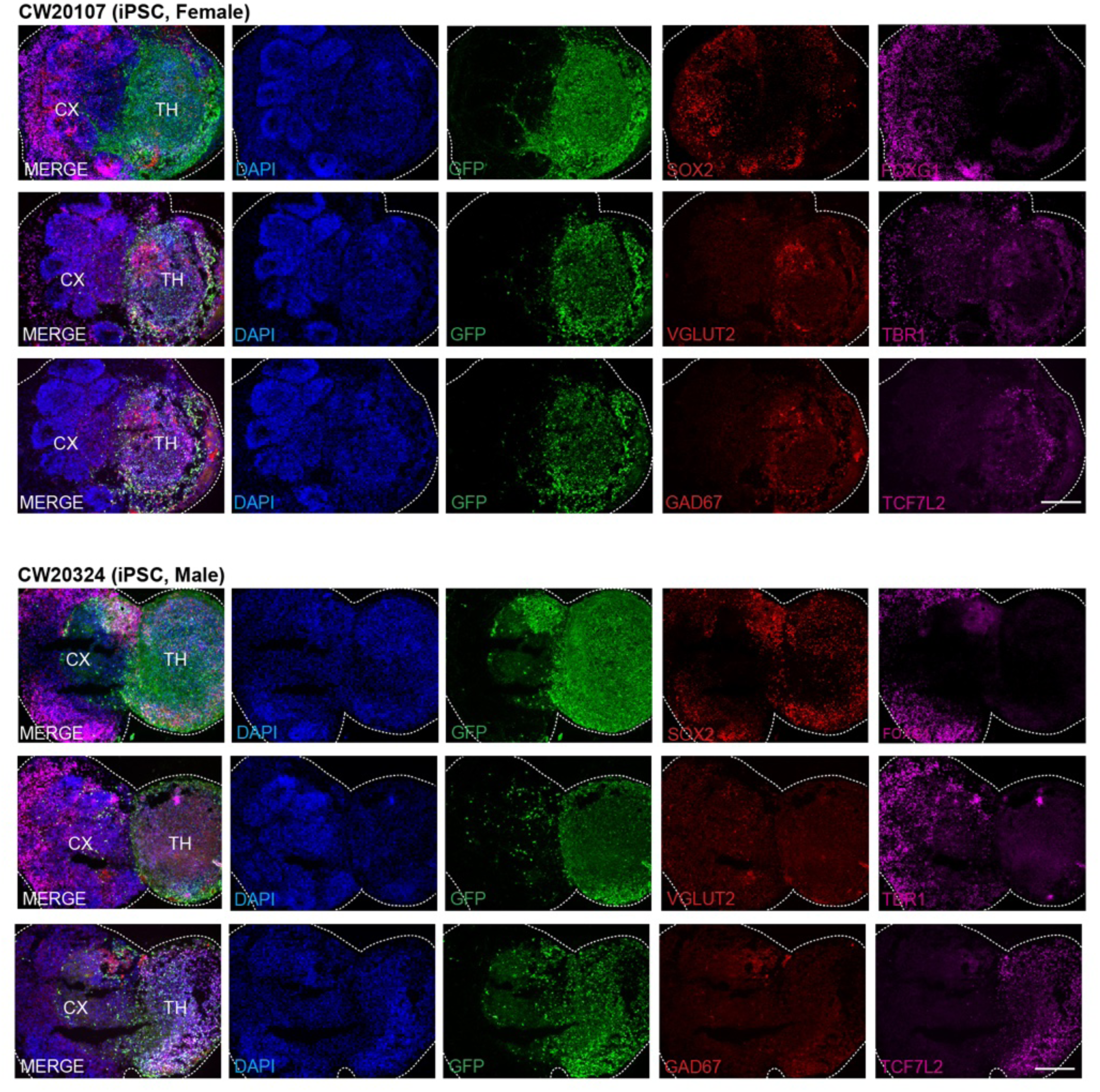
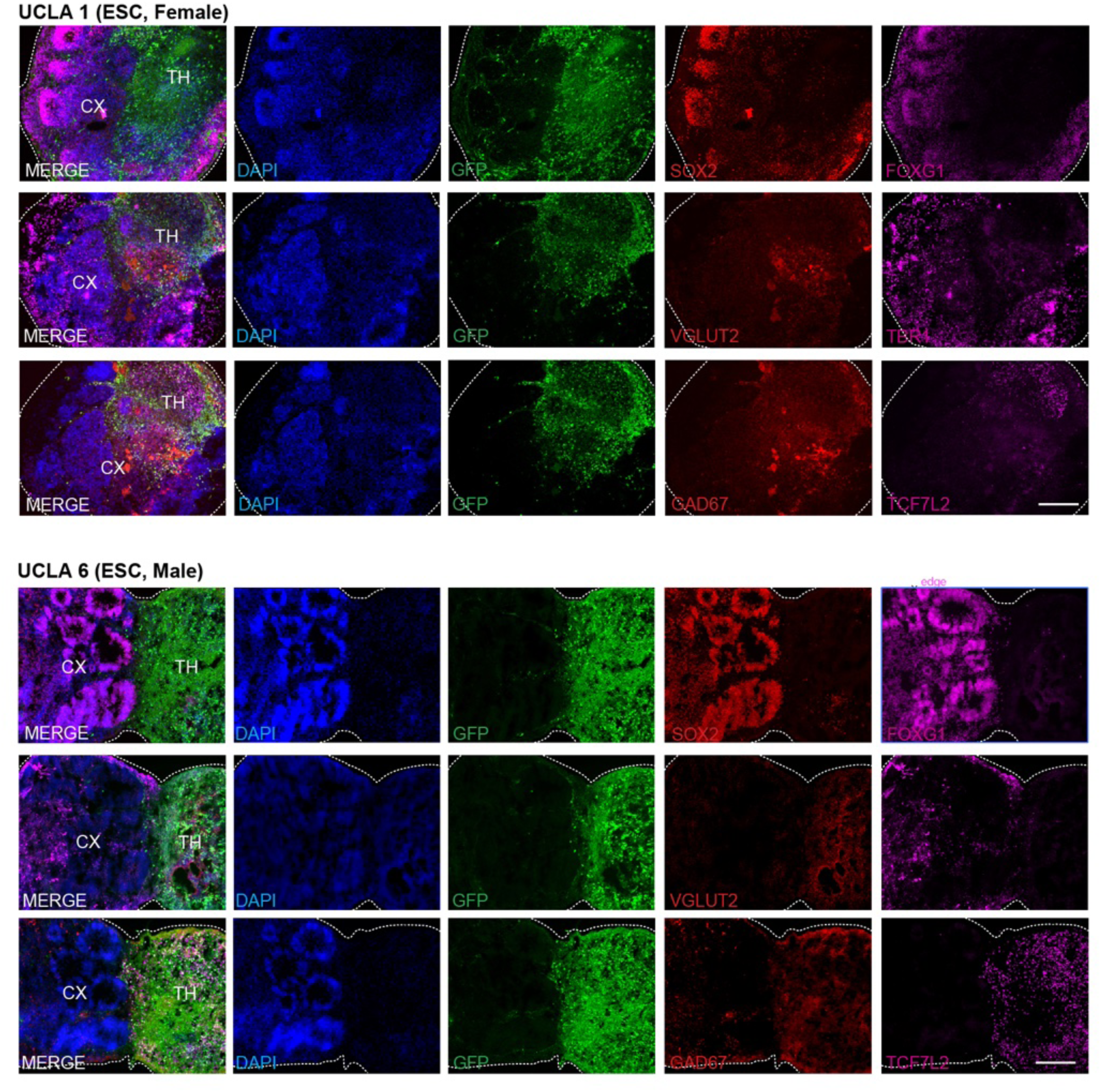
Cortical and thalamic identity markers are maintained in thalamocortical assembloids generated from human female and male ESC and iPSC lines. TC assembloids cultured in 50% cortical + 50% thalamic media were immunostained at week 8 to verify both sides maintained canonical markers for either cortical or thalamic fate post-fusion. Assembloid cell identities were identified by counterstaining for GFP, where GFP+ cells indicate thalamic regions and GFP-cells indicate cortical regions. Accordingly, GFP+ areas were positive for thalamus-enriched markers VGLUT2 (glutamatergic), GAD67 (GABAergic), and TCF7L2 (pan-thalamic). Conversely, GFP-areas expressed cortex-enriched markers FOXG1 (cortical progenitors) and TBR1 (cortical layer 6). As expected, both thalamic and cortical areas also contained cells positive for radial glia marker SOX2. Panels depict cell type marker panels for 4 cell lines from different genetic backgrounds: hiPSCs CW20107 (F) & CW20324 (M), and hESCs UCLA 1 (F) & UCLA 6 (M). Panels for the fifth cell line, UCLA 2, is depicted in main Figure 1. For each line, 3 - 4 technical replicates were examined. Scale bar = 250 μm.

**Supplementary Figure 3.**
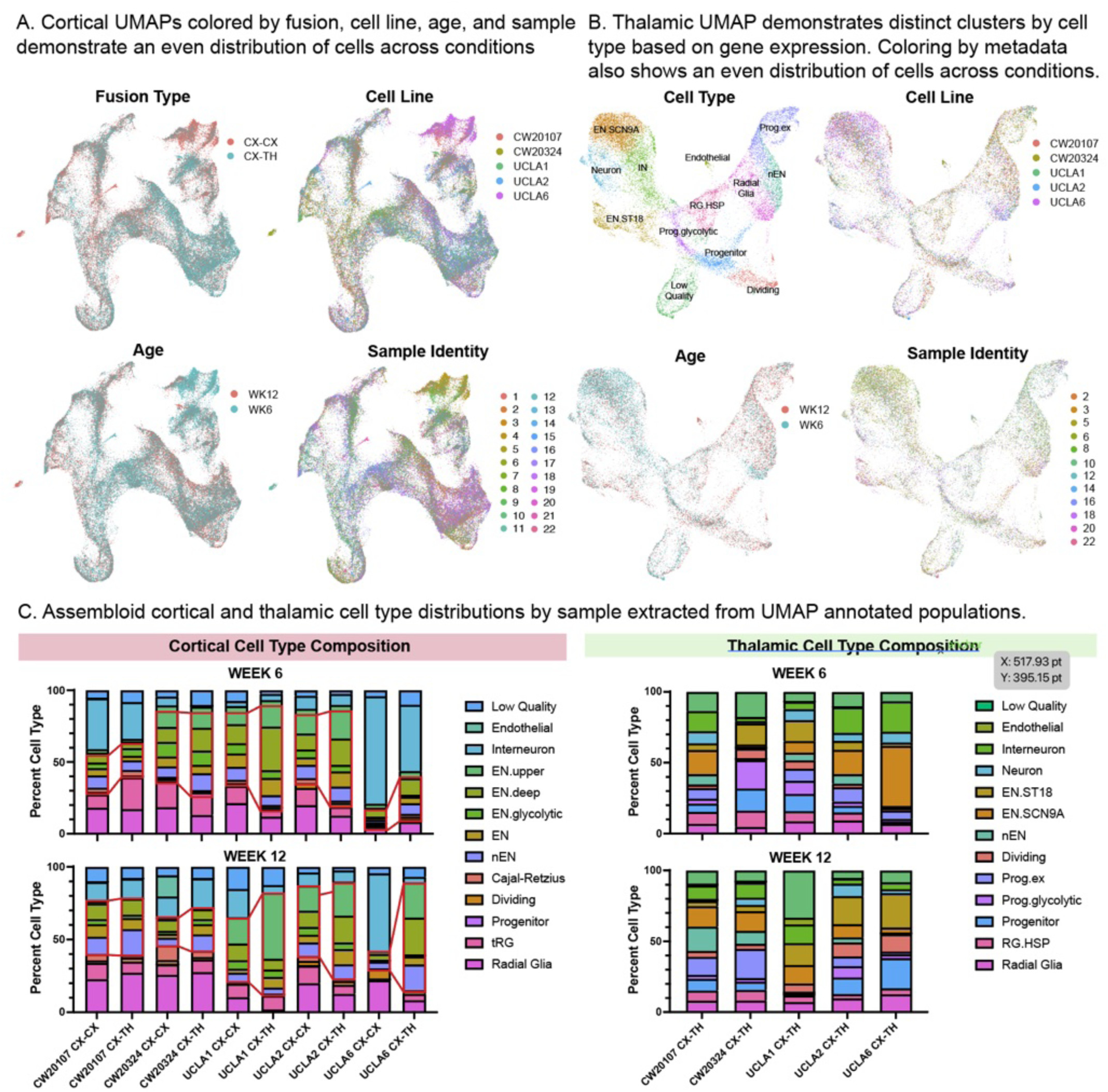
Analysis of cortical and thalamic snRNA-seq data validate an even distribution of cells across various sample conditions. D) Cortical cells from both CX-TH and control CX-CX assembloids were clustered using Seurat v5 and colored by various categories: assembloid type, cell line, age, and sample identity, as described in the metadata. The color-coded UMAPs demonstrate an even distribution of cells from each condition across cell type clusters (shown in Fig 1D). Each cluster has a representation of cells across assembloid type (CX-CX, CX-TH), cell line (CW20107, CW20324, UCLA 1, UCLA 2, UCLA 6), age (week 6, week 12), and sample identity (1–22), although some clusters may be enriched for certain conditions over others. E) Thalamic cells from CX-TH assembloids were also clustered and annotated using Seurat v5. The Cell Type UMAP depicts the annotated clusters’ names. Similar to cortex, the color-coded UMAPs demonstrate that there is an even distribution of cells from each condition across cell type clusters. Each individual cluster contains representation of each cell line (CW20107, CW20324, UCLA 1, UCLA 2, UCLA 6), age (week 6, week 12), and sample identity (1–22). F) The percentage of each cell type, segregated by cell line, was extracted from the cortical and thalamic UMAPs, respectively. Bar graphs depict the percentage of each annotated cell type in the cortical cells (left) and thalamic cells (right) across weeks 6 (top) and 12 (bottom). Red boxes highlight increases in excitatory neuron populations when comparing control (CX-CX) to thalamic-fused CX (CX-TH), which is observed across cell lines at both time points.

**Supplementary Figure 4.**
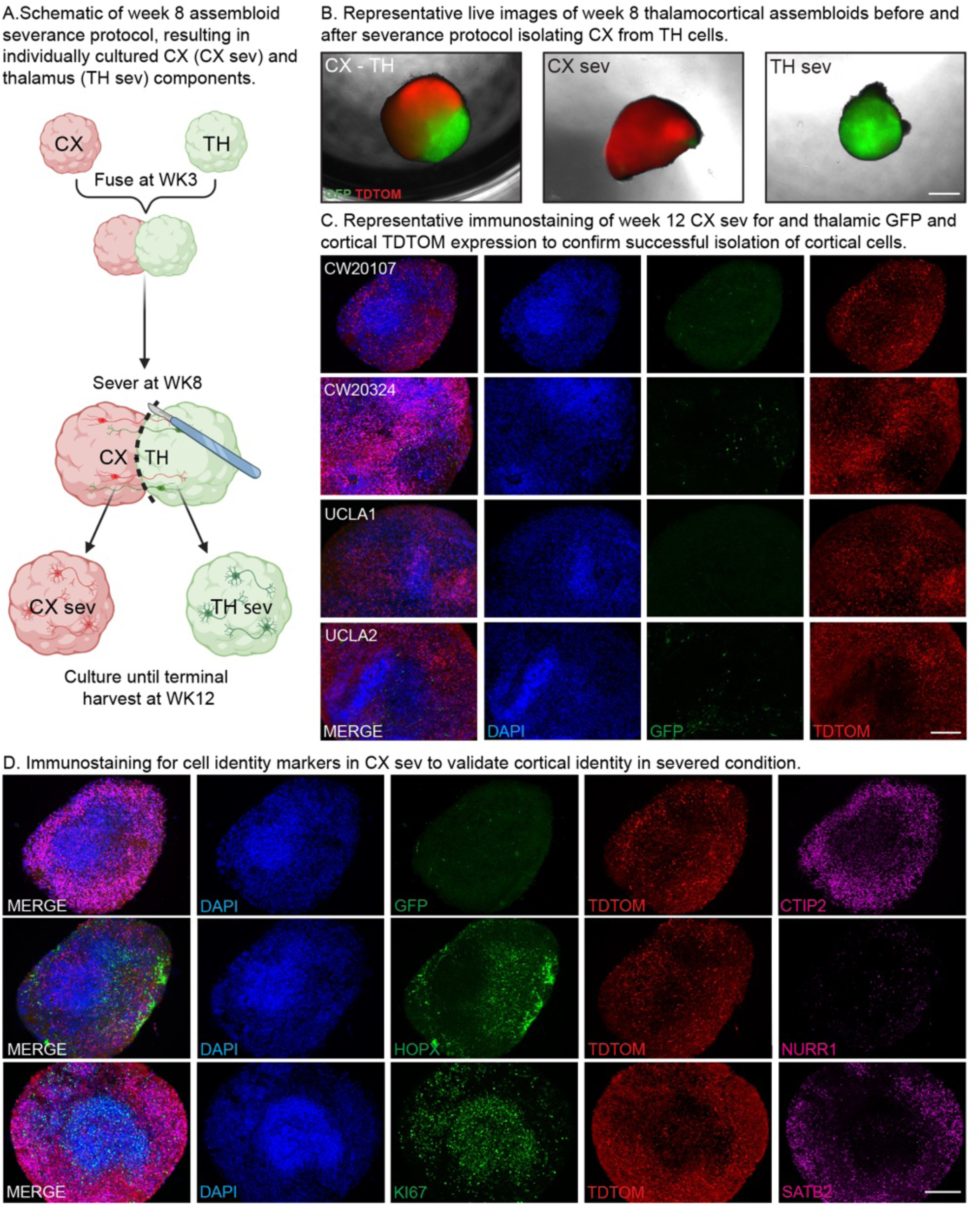
Severance of thalamocortical assembloids at week 8 enable assays of cortical cell types following the removal of thalamic input during mid-neurogenesis. A) Schema of TC assembloid severance protocol. Thalamic and cortical organoids were fused at week 3, then TC assembloids were severed with a scalpel at week 8, after which cortical (CX sev) and thalamic (TH sev) halves were cultured separately in 50/50 media. At week 12, both severed and intact assembloids were fixed for immunofluorescence staining. Generated using BioRender. B) Representative brightfield live images of week 8 TC assembloids before and after severance protocol isolating CX from TH cells. Left image depicts the intact CX-TH assembloid prior to severance, with red labeling tdTomato+ cortical and green labeling GFP+ thalamic cells. Middle image shows the cortical half of the assembloid post severance (CX sev), while the right image shows the thalamic half of the assembloid post-severance (TH sev). Scale bar = 500 μm. C) Immunofluorescence images across 4 cell lines demonstrating near complete depletion of GFP+ thalamic cells and afferents from CX sev condition at week 12 timepoint. TdTomato staining demarcates virally labeled cortical cells. Scale bar = 250 μm. D) Week 12 CX sev organoids were immunostained for cell identity markers to ensure retention of cortical identity in the severed condition. GFP (green, 1st row) and tdTomato (red, rows 1-3) label thalamic and cortical cells, respectively. CTIP2 (magenta, row 1) demarcates deep layer neurons, while SATB2 (magenta, row 3) demarcates upper layer neurons. HOPX (green, row 2) labels oRG and KI67 (green, row 3) labels dividing cells (scale bar = 250 μm).

**Supplementary Figure 5.**
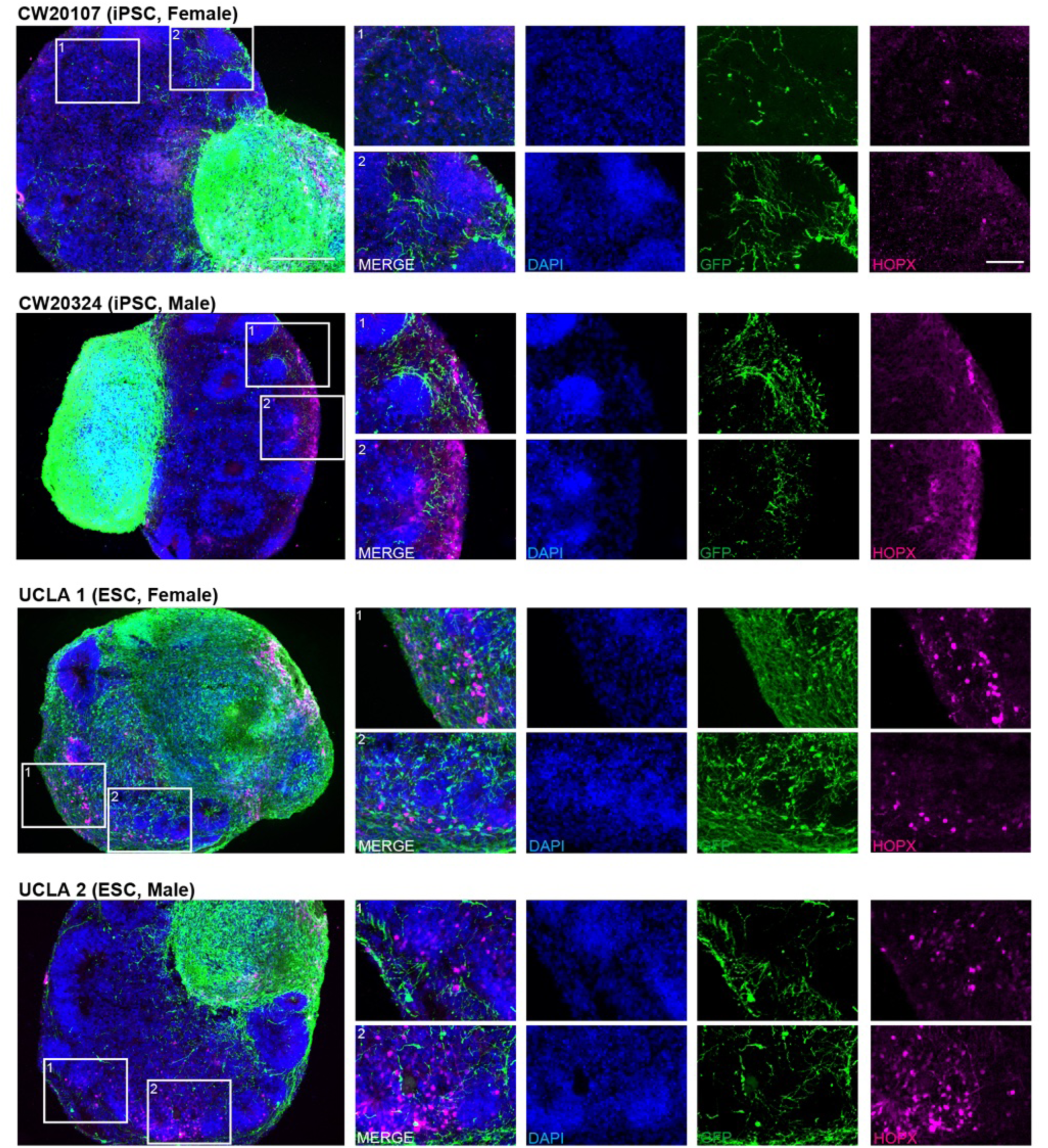
GFP+ thalamocortical afferents display targeted pathfinding around cortical organoid rosettes and colocalize with HOPX+ cortical outer radial glia as early as week 6 across lines. TC assembloids across five cell lines were immunostained to validate TCA projections into cortex. Assembloids were sectioned and stained for GFP+ thalamic afferents and HOPX+ (magenta) oRG (left, scale bar = 250 μm). Across both iPSCs and ESCs, TCAs are observed to infiltrate cortex and largely navigate around rosettes. Insets for each cell line highlight HOPX-dense areas where TCAs are found in close contact with cortical oRG (scale bar = 50 μm). Fifth cell line, UCLA 6, pictured in Figure 3A.

**Supplementary Figure 6.**
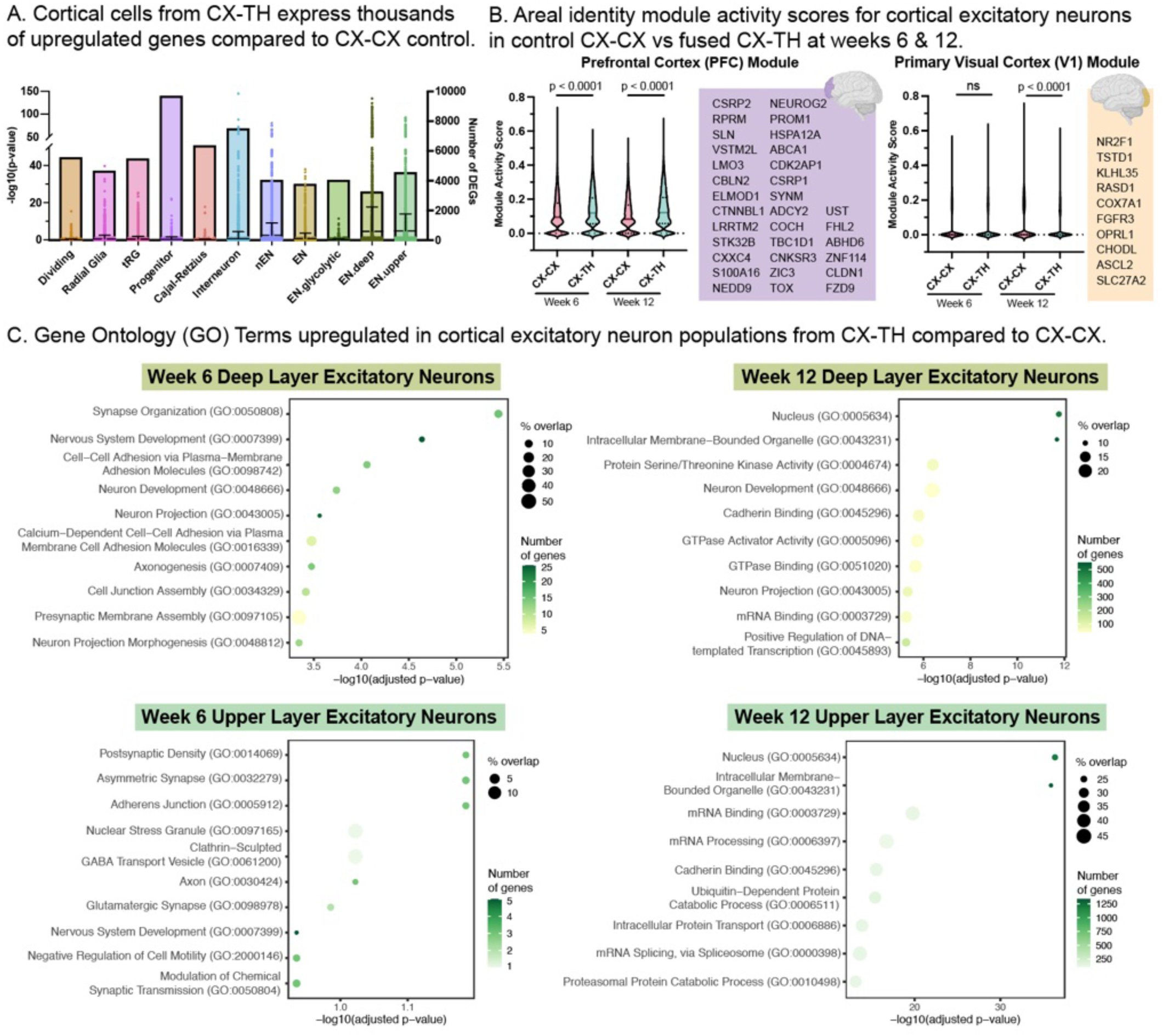
Differential gene expression analyses demonstrate transcriptional changes across cortical cell types from CX-TH compared to CX-CX control assembloids. A) Upregulated genes in cortical cell types from fused (CX-TH) as compared to control (CX-CX) from both week 6 and 12 data, segregated by cell type. The total number of differentially expressed genes (DEGs) are depicted by the bars (right axis). Individual genes and their - log10(p-value) are displayed as dots with their standard deviation error bars (left axis). Across cell lines and types, there are thousands of differentially expressed genes with variable significance. While neuronal populations consistently have significant changes in gene expression as predicted, cortical radial glia unexpectedly have significantly upregulated genes across all 5 cell lines. B) Cortical excitatory neurons from CX-CX and CX-TH were scored for areal identity modules representing genes highly enriched in the prefrontal cortex (PFC) and the primary visual cortex (V1). Results demonstrate that at weeks 6 and 12, cortical excitatory neurons fused with thalamus exhibited a higher PFC module score compared to CX-CX controls (p < 0.0001, multiple t-test). Conversely, module scoring for V1 identity only yielded statistically significant differences at week 12, in which CX-TH cortical excitatory neurons demonstrated a modest decrease in V1 identity (p < 0.0001, multiple t-test). In violin plots, bold lines represent group means, while dotted lines demarcate interquartile ranges. Genes used for each area identity module are depicted in the accompanying purple and yellow boxes. C) Upregulated genes from cortical deep and upper layer excitatory neurons from CX-TH compared to CX-CX were analyzed using EnrichR to determine relevant gene ontology (GO) terms. GO terms are listed on the left y-axis and significance as a function of -log10(adjusted p-value) are depicted on the x-axis. Circle location marks significance, while circle color represents number of genes represented and circle size represents percent overlap with the published GO term gene list. Week 6 deep layer excitatory neurons from CX-TH were enriched for terms associated with synapse organization, neuron development, and axonogenesis (top left). Week 12 CX-TH deep layer excitatory neurons were similarly enriched for terms associated with neuron development, but were also significantly associated with enzyme activity and transcription (top right). Similarly, week 6 upper layer excitatory neurons were associated with synaptic transmission GO terms (bottom left). Week 12 upper layer excitatory neurons showed the greatest levels of significance and gene overlap, with GO terms enriched in CX-TH related to mRNA processing and cell metabolism (bottom right).

**Supplementary Figure 7.**
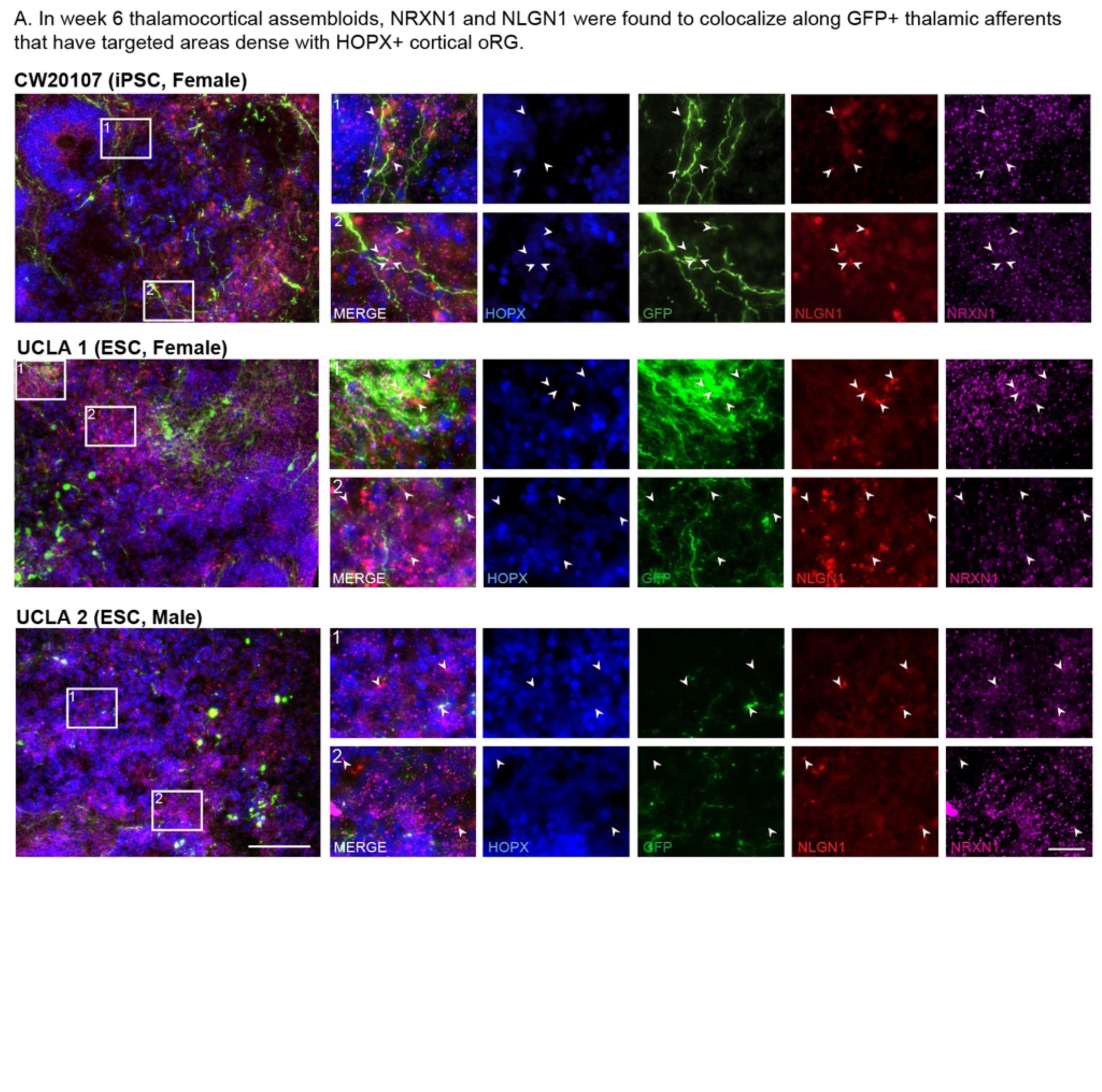
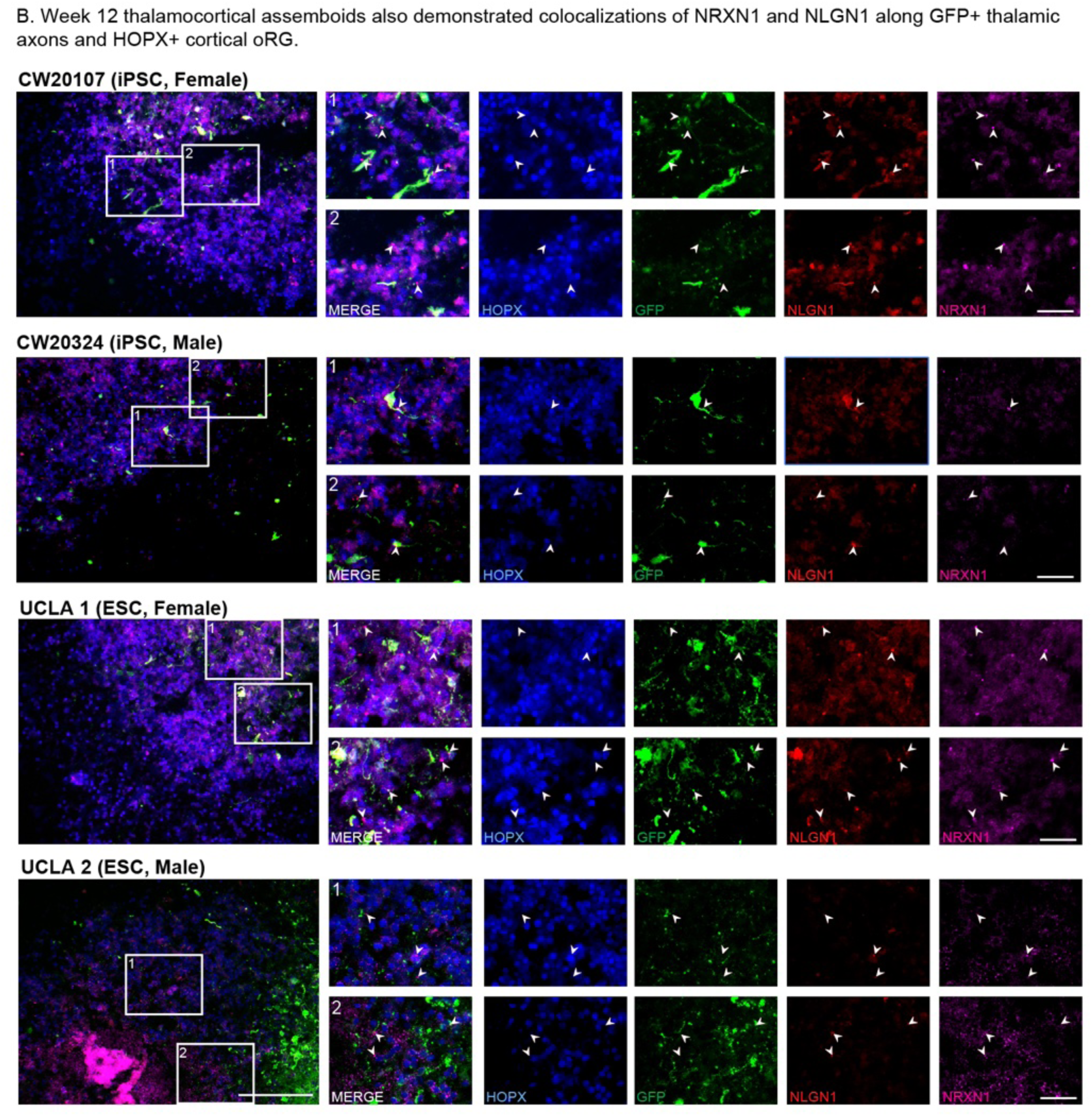
NRXN1 and NLGN1 are found to colocalize along GFP+ thalamic afferents targeting HOPX+ cortical radial glia in TC assembloids generated across cell lines. A) Panels depict representative images of the cortical half of week 6 TC assembloids across iPSC and ESC lines of varying genetic backgrounds. Staining shows areas where punctate proteins NRXN1 (magenta) and NLGN1 (red) are found to colocalize along GFP+ (green) thalamic afferents making contact with cortical HOPX+ (blue) oRG. 10X images are shown at left (scale bar = 250 μm), while insets depict zoomed in regions where NRXN1-NLGN1 colocalization is shown via white arrows (scale bar = 50 μm). Panel for fourth cell line, CW20324, is shown in main Fig 3. B) Images depict representative images of the cortical half of week 12 TC assembloids generated across the 4 main cell lines. Immunofluorescence staining depicts areas where GFP+ thalamic afferents (green) and HOPX+ oRG (blue) make physical contacts mediated by the colocalization of NRXN1 (magenta) and NLGN1 (red). 20X images are shown at left (scale bar = 125 μm), while insets depict zoomed in regions where NRXN1-NLGN1 colocalization is highlighted with white arrows (scale bar = 25 μm).

**Supplementary Figure 8.**
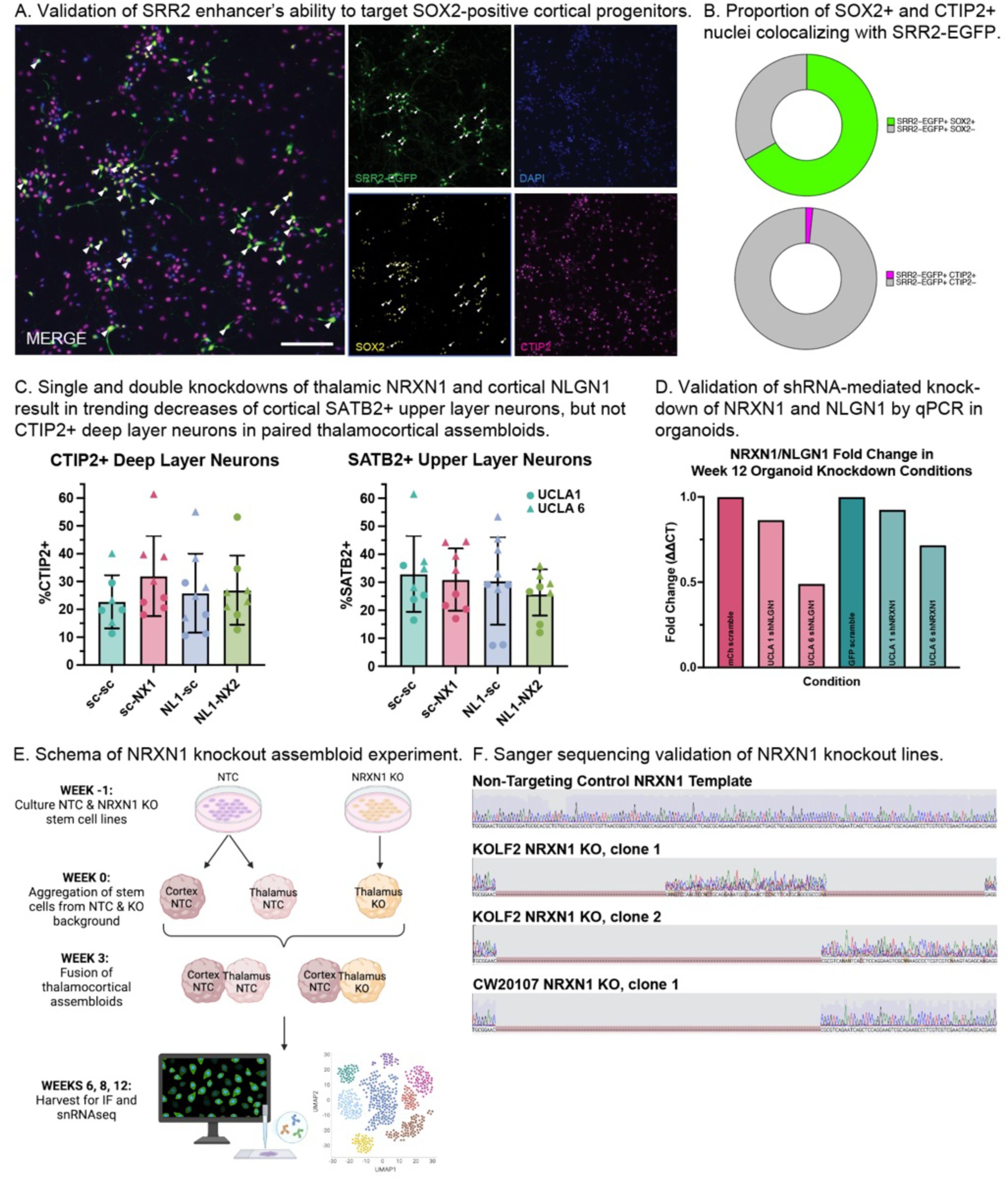
Validation of constructs and cell lines used in transcellular and NRXN1/NLGN1 perturbations *in vitro*. A) To validate the ability of SRR2-GFP virus to mark SOX2+ cortical progenitors, dissociated cortical organoid cell cultures were infected with pLEX-SRR2-GFP virus, then fixed and stained for GFP (green), SOX2 (yellow, radial glia), and CTIP2 (magenta, neurons). Images demonstrate that the vast majority of SRR2-GFP+ cells also colocalize with SOX2, a subset of which are labeled with white arrows (scale bar = 100 µM). B) Quantification of cell identity markers colocalizing with GFP demonstrates that 66.78% of SOX2-positive cells colocalize with SRR2-GFP+ cells, while only a small fraction (1.6%) of CTIP2 cells are SRR2-GFP+. This validates the ability of the SRR2-GFP virus to identify SOX2+ cortical progenitors *in vitro*. C) Short-hairpin RNAs (shRNAs) generated to target knockdown of thalamic NRXN1 and cortical NLGN1 in TC assembloids. Single- and double-knockdowns of thalamic NRXN1 and cortical NLGN1 were compared to scramble-only controls when quantifying cell types as marked by immunofluorescence markers for deep layer (CTIP2+) and upper layer neurons (SATB2+), analogous to analyses performed in Figures 1C and 2F. Quantification of these cell types demonstrated a trending, but non-significant decrease in cortical SATB2+ upper layer neurons, but not CTIP2+ deep layer neurons in paired CX-TH assembloids. This trend suggested that NRXN1/NLGN1 interactions may play a role in upper layer expansion and prompted the investigation of complete thalamic NRXN1 knockout models in Figure 5. D) Short-hairpin RNA (shRNA) viral constructs were validated by qPCR for their ability to knockdown NRXN1 and NLGN1 in week 12 thalamic and cortical organoids, respectively. E) Schematic depicting NRXN1 knockout assembloid experiment, in which stem cell lines with NRXN1 knockout (NRXN1 KO) were used to generate thalamic organoids. Non-targeting control (NTC) stem cell lines were used to generate both cortical and thalamic organoids. Assembloids were generated as described in Methods, and assembloids of NTC CX - NRXN1 KO TH were compared to those of NTC CX - NTC TH at weeks 6, 8, and 12. F) Sanger sequencing validates deletion of target DNA sequences in the NRXN1 locus in genetically edited cell lines (rows 2-4) as compared to non-targeting control intact NRXN1 template (row 1).

**Table S1.** Cell Line Information

**Table S2.** Viruses & Plasmids

**Table S3.** Antibodies

**Table S4.** snRNA-seq Analysis Metadata

